# simATAC: a single-cell ATAC-seq simulation framework

**DOI:** 10.1101/2020.08.14.251488

**Authors:** Zeinab Navidi, Lin Zhang, Bo Wang

## Abstract

Single-cell Assay for Transposase-Accessible Chromatin sequencing (scATAC-seq) identifies regulated chromatin accessibility modules at the single-cell resolution. Robust evaluation is critical to the development of scATAC-seq pipelines, which calls for reproducible datasets for benchmarking. We hereby present the simATAC framework, an R package that generates scATAC-seq count matrices that highly resemble real scATAC-seq datasets in library size, sparsity, and chromatin accessibility signals. simATAC deploys statistical models derived from analyzing 90 real scATAC-seq cell groups. simATAC provides a robust and systematic approach to generate in silico scATAC-seq samples with known cell labels for assessing analytical pipelines.

## Background

Single-cell sequencing has revolutionized and expedited our understanding of the structure and function of cells at unprecedented resolution. This technology resolves a fundamental limitation of bulk sequencing, which averages signals over a large number of cells resulting in obscured biological heterogeneity among individual cells [1]. The assay for transposase-accessible chromatin sequencing (ATAC-seq) measures chromatin’s openness, a proxy for the activity of DNA binding proteins [2–4]. Single-cell ATAC-seq (scATAC-seq) has opened up vast fields of applications, including extracting accessibility and co-accessibility patterns of genomic regions to identify cell-type-specific enhancers, chromatin heterogeneity, and transcription factor activities.

The rapid advancement of scATAC-seq technology has given rise to the development of computational tools for scATAC-seq data, and the integrative analysis of transcriptomic and epigenomic profiles [5–7]. Though simulated datasets with known cell labels have been the most common approach to benchmark the performance of analytical pipelines, there is no existing standard practice or simulation tool available to generate synthetic scATAC-seq datasets from real single-cell samples. Some previous studies generated in silico data by downsampling reads from bulk or existing scATAC-seq data, or deploying simple sampling algorithms [8–18]. However, these simulation methods were implemented as part of scATAC-seq analytical tool development and were usually incompletely documented, resulting in a lack of reproducibility. Further, due to the sparsity and noisy nature of scATAC-seq data, generating synthetic samples that closely resemble real datasets remains challenging.

Most scATAC-seq analytical pipelines consist of preprocessing (read quality control and alignment), core analysis (feature matrix generation) and downstream analyses (e.g. cell type clustering). A feature matrix summarizes the filtered reads from BAM files by counting the number of aligned reads that overlap within the defined genomic regions. The features represent a subset of genomic regions with specified genomic positions, nucleotide patterns, or biological functions [8, 19, 20]. A commonly used feature matrix for scATAC-seq is the peak-by-cell matrix that captures the highest accessibility signals from genomic regions (peaks) obtained from bulk ATAC-seq or single cells. However, a sufficient number of cells are required to identify such peaks, and consequently, the peak-by-cell feature matrix usually fails to recognize the rare cell type regulatory patterns [11]. Alternatively, the bin-by-cell feature matrix is generated by segmenting the whole genome into uniformly-sized non-overlapping “bins” and mapping the read counts to each bin [11]. Unlike the peak-by-cell matrices, the uniform segmentation of the genome does not screen out any genomic region, and thus has the potential to detect rare cell groups.

We hereby propose simATAC, a scATAC-seq simulation framework that generates simulated samples resembling real scATAC-seq data. Given a real scATAC-seq feature matrix as input, simATAC estimates the statistical parameters of the mapped read distributions by cell type, and generates a synthetic count array that captures the unique regulatory landscape of cells with similar biological characteristics. We demonstrate that the synthetic samples generated by simATAC highly resemble real scATAC-seq datasets in library size, sparsity (proportion of zero entries), and averaged chromatin accessibility signals.

## Results

### simATAC framework

simATAC deploys statistical distributions to model the properties of a bin-by-cell count matrix for a group of cells with similar biological characteristics. The main modelling parameters include read coverage of cells (library size), non-zero cell proportion in each bin, and the average of read counts per bin (bin mean). Bin-by-cell matrix quantifies the number of open chromatin read fragments overlapping with the fixed-length bins (5 kbp windows) across the whole genome. For each user-input real scATAC-seq dataset, simATAC performs two core simulation steps: (i) estimating the model parameters based on the input bin-by-cell matrix, including the library sizes of the cells, the non-zero cell proportion and the read count average of each bin; (ii) generating a bin-by-cell matrix that resembles the original input scATAC-seq data by sampling from Gaussian mixture and polynomial models with the estimated parameters. simATAC outputs a count matrix as a SingleCellExperiment (SCE) object [21], with additional options to convert it to other formats of feature matrices. Figure 1 summarizes the simulation architecture of simATAC. We discuss the statistical modelling of simATAC in the next sections and Methods.

**Figure 1.**
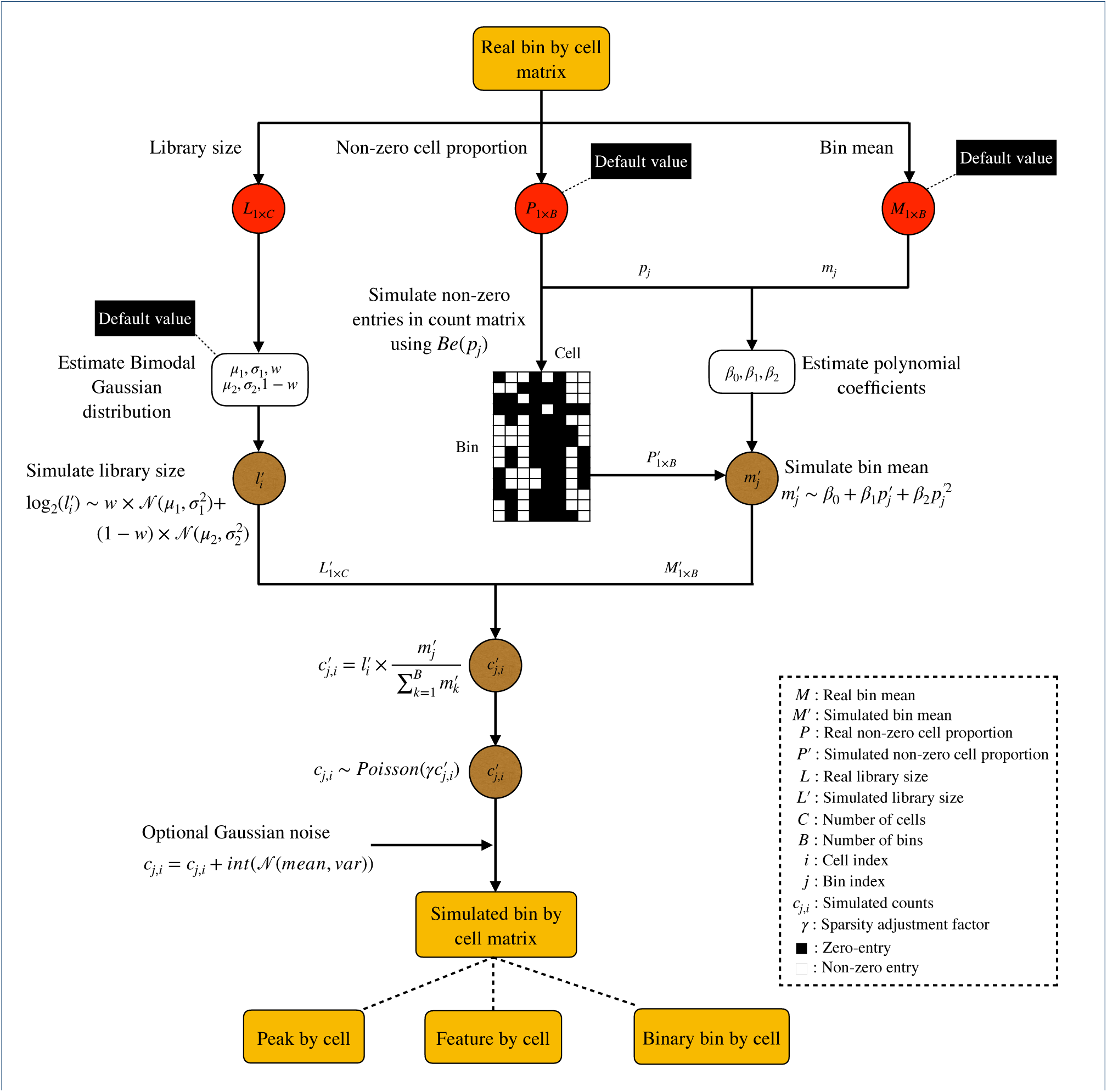
The simATAC simulation framework. The red circles represent values directly extracted from the user-input bin-by-cell matrix, the white squares represent estimated parameters, and the brown circles represent the simulated values. simATAC initially estimates the library sizes, the non-zero cell proportions, and the bin means from the input cell group (including cells having similar biological characteristics). simATAC generates the library sizes with a Gaussian mixture distribution, the zero and non-zero status with a Bernoulli distribution, and the bin means with a polynomial regression model linking to the non-zero cell proportion. The synthetic counts are sampled from a Poisson distribution whose mean is a factor of the cell library size adjusted by the bin mean. simATAC offers optional sparsity adjustment factor γ and noise parameters to adjust the sparsity and noise level of the synthetic counts generated from the Poisson distribution. The simATAC framework simulates synthetic scATAC-seq data using the default values for all the parameters if no user-input real data or parameters are given.

### Library size

Library size refers to the number of aligned reads per cell. simATAC models cells’ log-transformed library sizes through a Gaussian mixture model (GMM) with two components whose parameters are estimated based on the user-input real scATAC-seq data. With the estimated parameters, simATAC randomly samples library sizes of *C* single cells based on

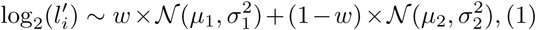

where 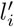 denotes simulated library size for the *i*^th^ simulated single-cell. See Table 1 for the detailed definition of the Gaussian mixture model parameters.

**Table 1.** Input parameters to the simATAC simulation step.

| Parameter | Symbol | Description |
| --- | --- | --- |
| Library size mean | $\mu_1, \mu_2$ | The estimated means of two Gaussian modals of library size. |
| Library size standard deviation | $\sigma_1, \sigma_2$ | The estimated standard deviations of two Gaussian modals of library size. |
| Library size weight | $w$ | The estimated weight parameter of the first Gaussian modal. |
| Non-zero cell proportion | $P$ | The proportion of non-zero cells in real bin-by-cell matrix. |
| Polynomial coefficient | $\beta$ | The estimated coefficients of polynomial model fitted to the relation between bin non-zero cell proportions and bin means. |

Previous studies have shown that the library sizes of cells significantly affect the identification of cell types [11–13, 15, 17]. Higher sequencing coverage usually results in a larger library size, thus providing more accurate chromatin accessibility information. Since the library sizes of scATAC-seq usually vary across different experiments, simATAC offers users the flexibility to adjust the library sizes from low to high coverage based on their needs.

### Bin non-zero cell proportion

Sparsity is the inherent nature of scATAC-seq data [11, 16, 17], which results in a large proportion of zero entries in the bin-by-cell matrix. Let *M*_*j,i*_ denote the number of reads that fall into the bin *j* of cell *i* for *B* bins. If *M*_*j,i*_ *>* 0, it is considered as a non-zero entry. The number of cells with non-zero entries within a bin is associated with the chromatin accessibility in the corresponding genomic region. Based on the user-input real scATAC-seq bin-by-cell matrix, simATAC first estimates the proportion of cells with non-zero entries for the *j*^th^ bin, *p*_*j*_, and then determines whether an entry in the simulated count matrix is zero or not based on a Bernoulli distribution,

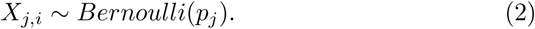

If *X*_*j,i*_ = 1, the read count of cell *i* at bin *j* is non-zero, i.e. *M*_*j,i*_ *>* 0. If *X*_*j,i*_ = 0, the read count of cell *i* at bin *j* is set to zero, i.e. *M*_*j,i*_ = 0. The non-zero cell proportion of bin *j*, 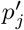, of the simulated bin-by-cell matrix is then defined as

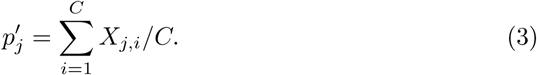

### Bin mean

The extent of genome accessibility leads to the variations in the number of sequenced reads falling into the fixed-length bins, and consequently, the variations in the average of the reads in each bin. More accessible regions potentially have larger bin means, that is, more cells with non-zero entries are mapped to that region. Based on the modelling scATAC-seq datasets, we observed a polynomial regression relationship between the non-zero cell proportions and the bin means for each cell group. simATAC simulates the average of the read counts at bin *j*, 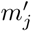, by

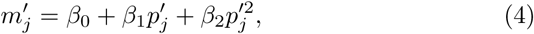

where *β*_0_, *β*_1_ and *β*_2_ are estimated based on the input real scATAC-seq dataset.

### Bin-by-cell count matrix

simATAC generates the final count of cell *i* at bin *j, c*_*j,i*_, using a Poisson distribution with 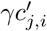 as the mean parameter, where 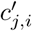 is the library size of cell *i* scaled by the bin mean 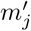 at bin *j*, and *γ* denotes the sparsity adjustment factor with a default value of 1. When *γ <* 1, the simulated scATAC-seq data tend to be more sparse, and vice versa.

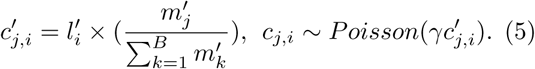

To reproduce the high noise level of scATAC-seq data, simATAC offers an optional step to include additional noise to the final simulated counts by

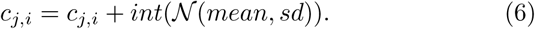

High noise level blurs the difference in the read distributions between different cell types, which mimics real sequencing artifacts. However, as the noise level increases, the distribution of the library sizes and the sparsity of the simulated data may differ from the input real scATAC-seq data. The default setting of simATAC omits the optional adding noise step, but leaves users the flexibility to set their desirable noise level.

simATAC outputs the final simulated bin-by-cell matrix as a SCE object, a container for single-cell genomics data, from the SingleCellExperiment R package [21].

### Simulating feature matrices in other formats

simATAC can also generate synthetic scATAC-seq data in other formats of feature matrices. The default bin-by-cell output can be converted to a peak-by-cell matrix by filtering bins enriched in aligned reads (top bins by measuring the bin means). simATAC generates a synthetic peak-by-cell count matrix with the “simATACgetPeakByCell” function, given a simulated bin-by-cell matrix by simATAC and a user-specified number of peak bins. simATAC also offers additional functionality to extract other feature matrices given a user-input list of regions (in a BED format) with the “simATACgetFeatureByCell” function. Another commonly used feature matrix format is the binary version, which is provided by “simATACgetBinary” function.

### Evaluation

In this section, we demonstrate the resemblance of the simulated samples generated by simATAC to the input real scATAC-seq datasets. The simulated samples are compared to the real samples on the distributions of library size, sparsity and bin means. We also evaluate the clustering performance of the simulated matrices. The evaluations are performed on each cell group (or cell type) from the annotated benchmark scATAC-seq datasets, Buenrostro2018 [22], Cusanovich2018 [23], and PBMCs [24], representing a wide range of platforms, cell types, and species. See Methods and Additional file 1: Table S1 for the description of cell types and platforms of these datasets.

### Statistical evaluation

With each of the three real scATAC-seq datasets as input, we simulate bin-by-cell matrices for each cell group with the same number of cells as in the real datasets. We then compare the distribution of library size, bin means, and sparsity of the simulated datasets to the real samples, by cell group. We present four cell groups from each benchmark dataset to demonstrate the similarity.

Figure 2 depicts the library size distributions of the scATAC-seq data simulated by simATAC using 12 cell groups from the benchmark datasets as input. The library size distributions of the simulated data highly resemble those of the real datasets. See Additional file 1: Figure S1 for the complete comparison between the real and simulated data for all the cell groups in the three benchmark datasets.

**Figure 2.**
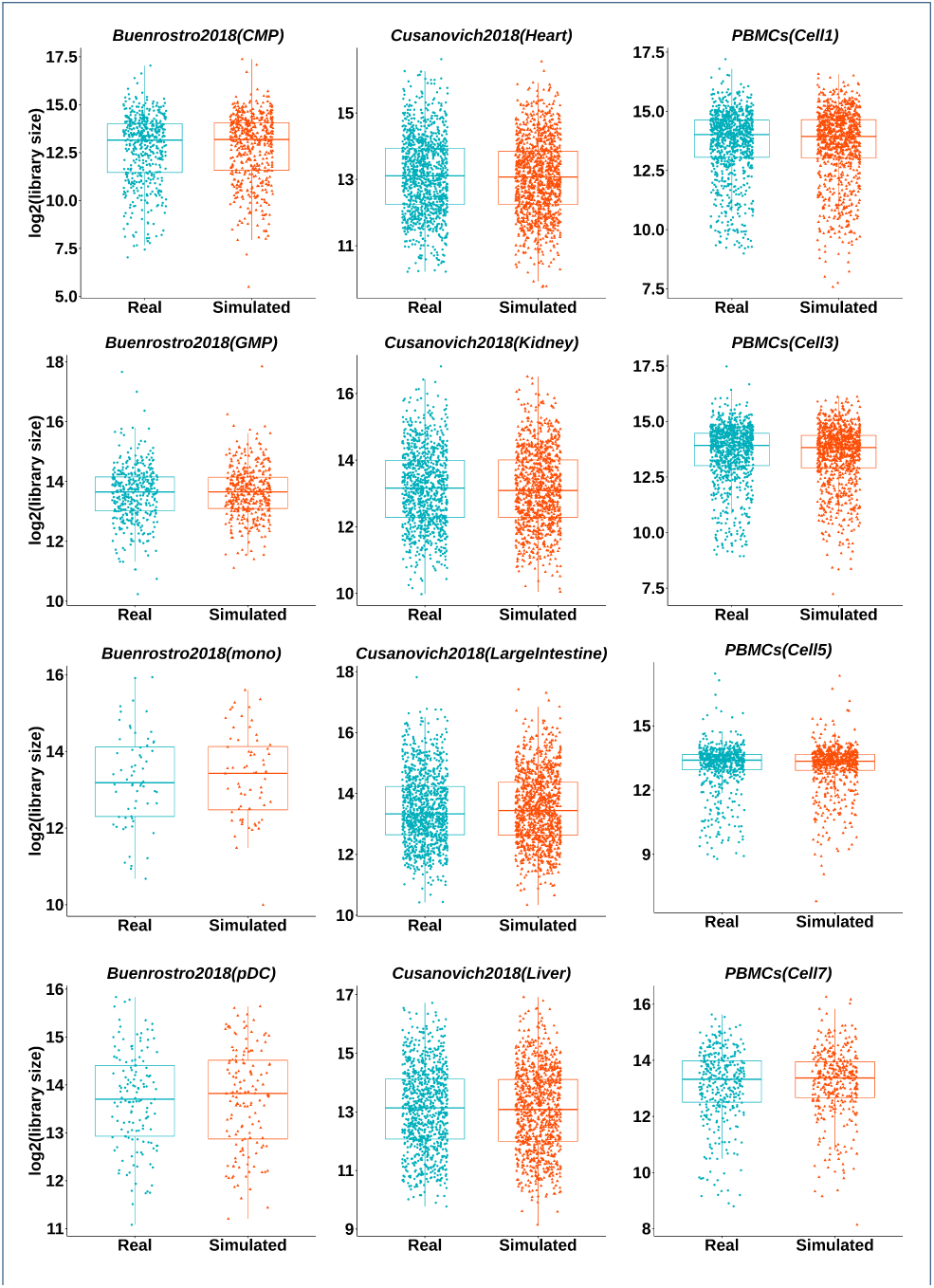
Comparison of the library size distribution. Library size box plots of the simulated (in orange) and real (in green) scATAC-seq data are illustrated for the three benchmark datasets: Buenrostro2018, Cusanovich2018, and PBMCs. The library size distributions of the synthetic samples (without additional Gaussian noise) closely resemble those of the real samples.

simATAC synthesized bin-by-cell matrices preserve the accessibility of genomic regions with its input real scATAC-seq dataset. Table 2 summarizes the Pearson correlation of the bin means and non-zero cell proportions between the simulated and the input real scATAC-seq datasets. The high correlations demonstrate that simATAC retains the genomic region accessibility characteristics of the input real data. See Additional file 1: Table S2 for the complete comparison for all cell groups from the benchmark datasets. All the reported Pearson correlations are averaged over 20 simulation runs.

**Table 2.** Pearson correlation between the simulated and real samples’ bin means and the non-zero cell proportion of bins.

| Cell type | Correlation |  |
| --- | --- | --- |
|  | Bin mean | Non-zero cell proportion |
| *Buenrostro2018* |  |  |
| CMP | 0.96 | 0.98 |
| GMP | 0.98 | 0.99 |
| pDC | 0.97 | 0.97 |
| mono | 0.92 | 0.94 |
| *Cusanovich2018* |  |  |
| Heart | 0.95 | 0.96 |
| Kidney | 0.97 | 0.97 |
| LargeIntestine | 0.93 | 0.93 |
| Liver | 0.98 | 0.98 |
| *PBMCs* |  |  |
| Cell1 | 0.99 | 1 |
| Cell3 | 0.99 | 1 |
| Cell5 | 0.98 | 0.99 |
| Cell7 | 0.99 | 0.99 |

Figure 3 illustrates the sparsity of the simulated bin-by-cell matrices in 12 cell groups of the benchmark datasets, which demonstrates that the synthetic samples generated by simATAC retains the sparsity of the real samples across bins and cells. See Additional file 1: Figures S2-S3 for other cell groups comparison. To demonstrate the bin sparsity resemblance of the simulated to the real data, we provide the bin sparsity QQ-plots of 12 benchmark sample groups for sparsity adjustment factor *γ* = {0.8, 0.9, 1} in Additional file 1: Figures S4-S6.

**Figure 3.**
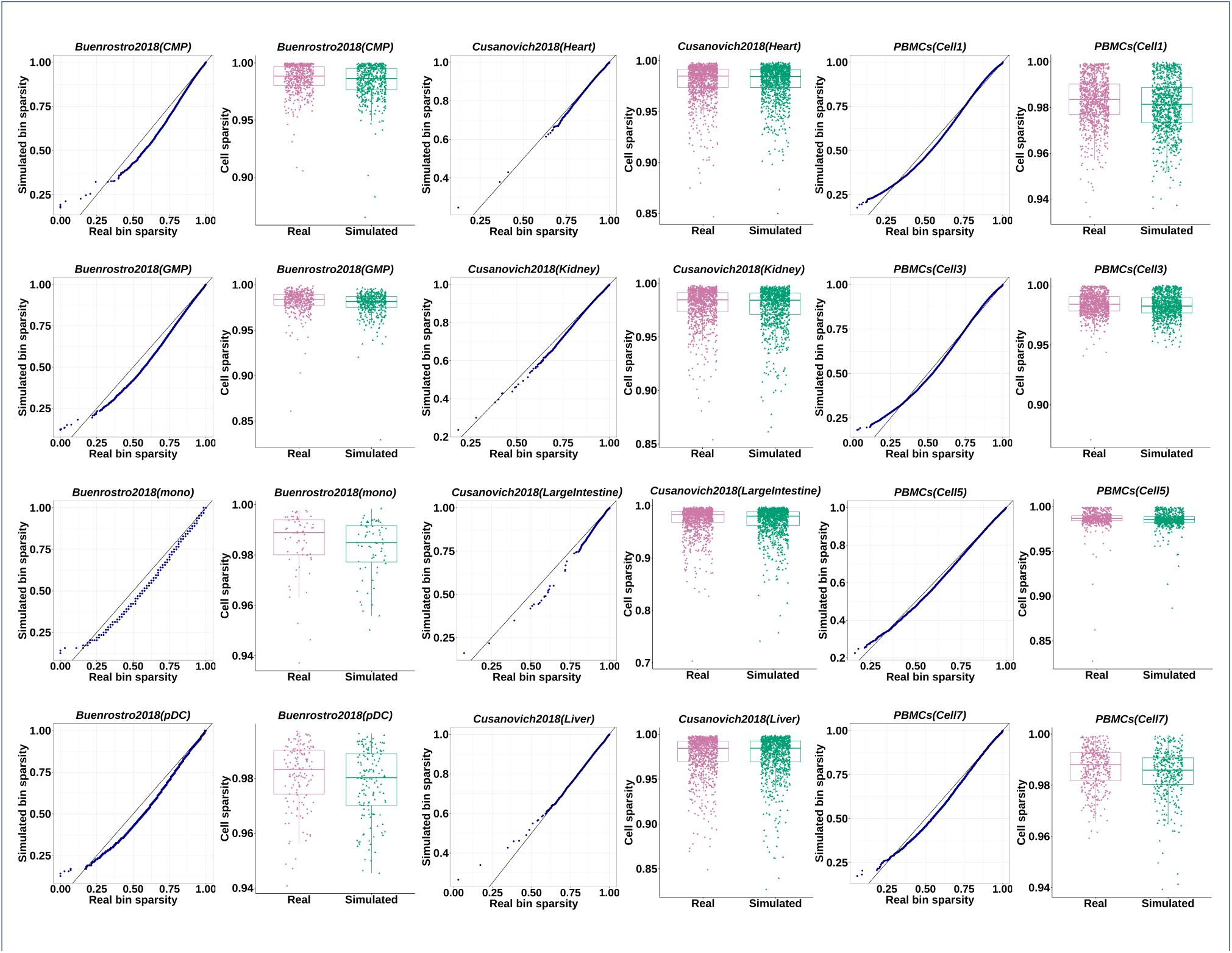
Comparison of the bin sparsity and cell sparsity distributions. Bin sparsity QQ-plots and cell sparsity box plots of the simulated (in green) and real (in purple) scATAC-seq data are illustrated for the three benchmark datasets: Buenrostro2018, Cusanovich2018, and PBMCs. The sparsity of the synthetic data generated by simATAC (without additional Gaussian noise) closely resembles that of the corresponding real scATAC-seq input data.

We further investigate the impact of the bin sparsity adjustment parameter *γ* on the simulated bin-by-cell matrices. We observe that mild changes in the sparsity adjustment factor do not significantly affect the distributions of cell sparsity, library sizes, or bin sparsity. See Additional file 1: Figure S7 for a comparison of these distributions under different sparsity adjustment factor values *γ* = {0.8, 0.9, 1, 1.1, 1.2} using the LMPP cell type from the Buenrostro2018 dataset.

We also report the median absolute deviation (MAD), mean absolute error (MAE), and root mean square error (RMSE) of the sorted library sizes, bin means and non-zero cell proportion of each bin between the real and the simulated datasets in Table 3. The reported metrics are averaged over 20 simulation runs. The small values of MAD, MAE, and RMSE suggest that the synthesized bin-by-cell matrices’ properties closely resemble those of the real input data.

**Table 3.** Median absolute deviation (MAD), mean absolute error (MAE), and root mean square error (RMSE) average for sorted real and sorted simulated library sizes, real and simulated bin means, and real and simulated non-zero cell proportions across all cell groups in the associated dataset. The reported values are the averages of these metrics the ± corresponding standard deviations based on 20 simulation runs. The unit of the values is 0.1%.

|  | Buenrostro2018 |  |  | Cusanovich2018 |  |  | PBMCs |  |  |
| --- | --- | --- | --- | --- | --- | --- | --- | --- | --- |
|  | MAD | MAE | RMSE | MAD | MAE | RMSE | MAD | MAE | RMSE |
| Non-zero cell proportion | 2.4 ± 2.8 | 6.5 ± 1.7 | 15.2 ± 3.8 | 4.0 ± 1.4 | 5.8 ± 1.7 | 8.6 ± 2.3 | 5.8 ± 9.1 | 9.0 ± 14 | 13.9 ± 18 |
| Bin mean | 2.4 ± 2.8 | 9.0 ± 2.5 | 531.9 ± 21 | 4.4 ± 1.6 | 6.6 ± 2.0 | 32.2 ± 16 | 7.2 ± 12 | 13.1 ± 20 | 57.8 ± 59 |
| Library size | 90 ± 46 | 111.6 ± 47 | 150.2 ± 60 | 50.0 ± 18 | 63.3 ± 17 | 88.2 ± 23 | 72.6 ± 38 | 107.8 ± 44 | 169 ± 45 |

The main functionality of simATAC is to simulate bin-by-cell scATAC-seq count matrices, yet it also offers additional functionalities such as converting the simulated bin-by-cell matrices to peak-by-cell feature matrices using the “simATACgetPeakByCell” function. We show that the extracted peak-by-cell matrices preserve the chromatin accessibility information of the real input data. We compare the clustering performance and peak overlaps of the simulated data with those extracted from two commonly used non-segmenting peak callers, MACS2 [25] and Genrich [26]. See Additional File 1: Figure S8, Tables S3-S4, and Note S2 for the detailed comparison.

### Clustering evaluation

The ability to cluster cells with similar biological characteristics is one of the major evaluation aspects of many scATAC-seq analytical tools. Many previous studies reported close to perfect cell clustering performance when evaluating scATAC-seq pipelines using simulated data [11, 12, 17], which is not ideal in comparing the performance across different computational tools.

We here show that the simulated data by simATAC produce realistic downstream analysis results. Further, if the data is simulated based on an input real scATAC-seq dataset, the clustering performance based on the simulated data closely resemble those based on the real input data. Table 4 summarizes the clustering metrics, normalized mutual information (NMI), adjusted mutual information (AMI), and adjusted Rand index (ARI) evaluated on the simulated datasets by simATAC. We adopt the SnapATAC graph-based clustering algorithm, and all the reported metrics are averaged over 20 simulation runs [11].

**Table 4.** Clustering evaluation results. The NMI, AMI, and ARI scores are the cell type clustering results using SnapATAC software. “Real-input data” refers to the clustering results using the input real scATAC-seq data. Metrics for simATAC’s simulated bin-by-cell matrices for different noise levels are also compared presented for each of the Buenrostro2018, Cusanovich2018 and PBMCs datasets.

| Mean | Standard deviation | NMI | AMI | ARI |
| --- | --- | --- | --- | --- |
| *Buenrostro2018* |  |  |  |  |
| 0 | 0 | 0.71 | 0.83 | 0.54 |
| -0.3 | 0.3 | 0.72 | 0.83 | 0.54 |
| -0.4 | 0.4 | 0.67 | 0.78 | 0.50 |
| Real-input data |  | 0.54 | 0.62 | 0.36 |
| *Cusanovich2018* |  |  |  |  |
| 0 | 0 | 0.96 | 0.97 | 0.94 |
| -0.3 | 0.3 | 0.79 | 0.85 | 0.69 |
| -0.4 | 0.4 | 0.62 | 0.71 | 0.40 |
| Real-input data |  | 0.41 | 0.46 | 0.22 |
| *PBMCs* |  |  |  |  |
| 0 | 0 | 0.78 | 0.87 | 0.71 |
| -0.3 | 0.3 | 0.61 | 0.73 | 0.47 |
| -0.4 | 0.4 | 0.56 | 0.67 | 0.44 |
| Real-input data |  | 0.45 | 0.54 | 0.28 |

We vary the Gaussian noise levels (mean = 0, SD = 0; mean = −0.3, SD = 0.3; mean = −0.4, SD = 0.4) added to the simulated data, representing no noise to high noise levels. We compare the clustering metrics using the simulated data with those using the input real scATAC-seq data, which is listed on the “Real-input data” row. The clustering results in Table 4 show that the simulated data by simATAC maintain the chromatin accessibility information of each cell group reasonably close to the real data. The clustering metrics of the synthetic data can get very close to the real data when adding Gaussian noise, and achieve realistic clustering performance in contrary to existing simulation methods. We further assess the distributions of the simulated counts under these three recommended noise levels in Additional file 1: Figures S9-S14 and Tables S5-S6, which suggests that there is a trade-off between the closeness in clustering performance and the closeness in the distributions of library size and the sparsity parameters between the simulated and real samples. We also note that the impact of Gaussian noise level on clustering metrics may vary across different cell types, and thus simATAC offers the flexibility for users to adjust their desirable noise level.

In Table 4, we assume the sparsity parameter *γ* = 1 when simulating all the data. We show in Additional file 1: Table S7 the impact of *γ* on the downstream clustering performance of the simulated data. With all other parameters fixed, the clustering results are generally consistent across different values of *γ*. Hence, we set the default value of *γ* to 1 in simATAC.

## Discussion and conclusions

The rapid development of scATAC-seq technology led to a surge of scATAC-seq analytical tools. However, the lack of systematic simulation frameworks hinders the consistent evaluation of the computational tools and reproducibility of the analytical results. To meet this need, we developed simATAC, a systematic scATAC-seq simulator that generates synthetic samples that closely resemble real scATAC-seq data. simATAC builds upon Gaussian mixture distribution to model cells’ library size, and polynomial regression model to represent the relationship between the average of bin counts and the non-zero cell proportion of bins. Moreover, simATAC grants users the flexibility to adjust parameters manually. simATAC generates a synthetic bin-by-cell matrix given a real scATACseq dataset as input. If there are no user-specified count matrix or parameters available, simATAC simulates samples using the default parameters derived from real scATAC-seq data. A list of estimated values for library size Gaussian mixture distribution and polynomial function parameters are provided in Additional file 2. simATAC also offers additional functions to transform the bin-by-cell matrix into feature matrices in other formats.

The statistical modelling framework of simATAC is built upon 90 real scATAC-seq cell groups from various sequencing platforms, species, and cell types. We demonstrated the distributions of the library sizes, bin means, and sparsity parameters of the simATAC synthetic datasets resembling those of the real input examples. simATAC offers additional options to modify the noise levels to mimic the artifacts in real scATAC-seq data and generate samples with various difficulty levels for downstream analyses assessment.

Quality control steps in most scATAC-seq pipelines apply filters on the raw feature matrices, such as removing cells with a library size less than a fixed threshold or filtering out regions based on the number of mapped reads. As the denoising thresholds may affect the downstream analyses, simATAC offers users the flexibility to manually set the quality control filtration thresholds on the simulated raw count matrix.

The feature matrices generated by simATAC cover regions spanning the whole genome without discarding the off-peak reads, enabling the identification of rare cell types in complex tissues. While the peak-by-cell matrix has been the common version of scATAC-seq feature matrix to be analyzed, recent studies challenged this strategy and proposed the bin-by-cell version for downstream analyses. Peak calling pipelines perform differently in defining accessible genomic regions based on the approach they deploy. Therefore, we believe genome binning is an optimal approach to simulate scATAC-seq data, providing a standard representation for samples from different sources.

simATAC generates count matrices using a 5 kbp bin window, and at this resolution, each human cell spans ∼600,000 bins. However, there is no standard agreement on the optimal bin size for all samples, and bin size selection induces a trade-off between the ability to capture chromatin accessibility signals and computational cost. We assessed the runtime of simATAC on a desktop workstation (Intel(R) Xeon(R) CPU @ 3.60GHz processor). Simulating 1,000 human cells at 5 kbp window size on average took 43 seconds in five simulation runs. See Additional file 1: Table S8 for the running time of all benchmark datasets.

To our best knowledge, simATAC is the first scATAC-seq simulator that directly simulates bin-by-cell count matrices that are reproducible and closely resemble real data. Though the availability of real scATAC-seq data has been increasing, real scATAC-seq data with annotated labels (“ground truth”) remains lacking. simATAC offers users the convenience to generate scATAC-seq data with known cell types and desirable number of single cells, yet closely resemble the real data. Further, simATAC provides users the flexibility to adjust the library size, bin sparsity and noise level of the simulated data. We envision that simATAC empowers users to develop scATAC-seq analytical tools effectively and reproducibly.

## Methods

### simATAC statistical modelling

We compiled and processed each of the 90 scATAC-seq cell groups as well as each of the eight datasets (considering all cell groups together) to model the library size parameter. We conducted the Kolmogorov-Smirnov test and the Chi-squared test to test the goodness of fit of the log-transformed library sizes to the Gaussian probability distribution, using the stats and fitdistrplus R packages [27, 28]. The *p*-values of the goodness of fitness tests showed that non-10xG samples generally follow a Gaussian distribution. Our preliminary statistical analysis of the 10xG scATAC-seq data showed that many of them are sampled from a mixture of probability distributions. We tested the null hypothesis if the 10xG samples’ library sizes are sampled from a unimodal probability distribution using Hartigan’s dip test from the diptest R package [29]. In most of the modelling datasets, the null hypothesis is rejected at a significance level of *α* = 0.05. Considering the probability density function and the cumulative distribution function plots, we modelled the log-transformed library sizes with a Gaussian mixture model with two modes and estimated the parameters using the mixtools R package [30]. Statistical parameters of the aforementioned tests for library size modelling are provided in Additional file 3.

We observed a significant difference in the distribution of library sizes between the real scATAC-seq data generated by the 10xG platform and other platforms. Library sizes of the non-10xG samples generally fit a unimodal Gaussian model, while those of the 10xG samples fit a bimodal GMM. All the statistical analyses results are provided in the Additional file 3. simATAC simulates the library size using a bi-modal GMM for samples from all platforms, and for non-10xG samples, the weight of the second Gaussian distribution can be set to zero.

To recover the scATAC-seq data sparsity, simATAC first assigns zero or non-zero labels to the bin-by-cell matrix using a Bernoulli distribution for each bin. The probability that a cell at bin *j* is non-zero is the estimated non-zero cell proportion at the corresponding bin of the input real scATAC-seq dataset.

Based on the 90 real scATAC-seq modelling cell groups, we observed a polynomial relationship between the non-zero cell proportions and the bin means in the normalized real bin-by-cell arrays. The input matrix is normalized by dividing primary counts by the cells’ library size and multiplying by the median of library sizes. The quadratic relation between bin means and non-zero cell proportions for the 12 sampled cell groups of benchmark datasets are provided in Additional file 1: Figure S15. simATAC estimates the regression parameters using the lm function from the stats package [27], and calculates bin means based on equation Note that the parameters in equation 4 are estimated by cell types/groups, as the chromatin accessibility patterns of different cell types vary biologically.

### Evaluation metrics

We assessed the simulated feature matrices by calculating the absolute differences between sorted real and sorted simulated library size vectors, real and simulated bin means, and non-zero cell proportion vectors of original and synthetic count matrices. The MAD, MAE, and RMSE of these vectors are computed by the following equations, where R are the real values and S are the simulated values:

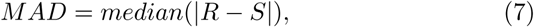

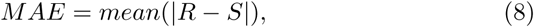

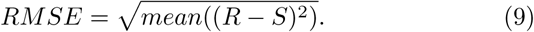

We used three metrics to evaluate the clustering performance, normalized mutual information (NMI), adjusted mutual information (AMI), and adjusted Rand index (ARI). Considering *gt* and *pred* as the ground truth and predicted labels, NMI is calculated using

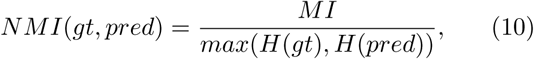

where *MI* = *MI*(*gt, pred*) is the mutual information (MI) between *gt* and *pred*, and *H* is the entropy.

AMI is defined as

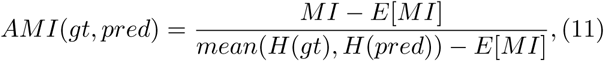

where *E*(·) is the expectation function. Using the same notations, ARI is defined as

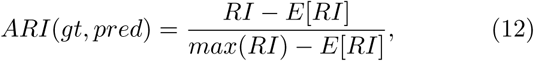

where the Rand index (RI) is a similarity measure between two lists of labels. See more details of NMI, AMI and ARI in Additional file 1: Note S2 [31].

### Availability of data and materials

We built the simATAC statistical model and estimated the default input parameters based on 90 cell groups from eight publicly available real scATAC-seq datasets from various platforms, including Fluidigm C1, 10x Genomics Chromium (10xG), single-cell combinatorial indexing ATAC-seq (sciATAC-seq), multi-index single-cell ATAC-seq (MI-ATAC), and single-nucleus ATAC-seq (snATAC-seq) to ensure a generalizable simulation framework. The datasets supporting the modelling and evaluation of this article are all available publicly. All datasets used in this study are available from GEO accessions: (1) 63 10xG sample groups from GSE129785 [32], (2) GSE99172 [8], (3) GSE74310 [33], (4) GSE65360 [34], (5) GSE68103 (GSM1647122) [35], (6) GSE68103 (GSM1647123) [35], (7) GSE112091 (series GSE112245) [36], and (8) GSE100033 (GSM2668124) [37].

The benchmark datasets used for evaluating simATAC framework are:

- The Buenrostro2018 dataset contains 1974 cells generated from the Fluidigm C1 platform. Samples are from 9 FACS-sorted cell populations from CD34+ human bone marrow, namely, hematopoietic stem cells (HSCs), multipotent progenitors (MPPs), lymphoid-primed multipotent progenitors (LMPPs), common myeloid progenitors (CMPs), granulocyte-macrophage progenitors (GMPs), megakaryocyte-erythrocyte progenitors (MEPs), common lymphoid progenitors (CLPs), plasmacytoid dendritic cells (pDCs), and monocytes (mono) [22].
- The Cusanovich2018 dataset is a subset of mouse tissues with 12178 cells from 13 different sources [17]. Sequenced cells are from bone marrow, cerebellum, heart, kidney, large intestine, liver, lung, prefrontal cortex, small intestine, spleen, testes, thymus, and whole brain, generated using a sciATAC-seq protocol [23].
- The PBMCs dataset is produced by the 10x Genomics Chromium (10xG) droplet-based platform and comprises 5335 cells from human peripheral blood mononuclear cells (PBMCs) [24]. There are no true cell type labels for PBMCs cells. However, we used 10x Genomics Cell Ranger ATAC’s [24, 38] clustering labels as ground truth and performed simulation on each cluster. Although existing labels are not perfect, we included this data to evaluate how simATAC mimics features of a group of cells with similar biological characteristics from a droplet-based platform [17].

All the aforementioned datasets are publicly available. The detailed information of all samples with cell groups and numbers are provided in the Additional file 1: Table S1. We also provide a table of abbreviations in Additional file 1: Table S9.

### Dataset pre-processing

The raw FASTQ or BAM files were downloaded from the links provided, and bin-by-cell matrices used in simATAC development were generated using the SnapTools (version: 1.4.6) [11, 39]. SnapTools is a Python module that pre-processes scATAC-seq data. SnapTools aligns raw FASTQ files to a reference genome using the Burrows-Wheeler aligner. Reads that were properly paired according to the SAM flag value, uniquely mapped with mapping quality *>* 30, and had a length less than 1,000 base pairs were filtered for further analyses. SnapTools groups the reads with the same barcode and removes PCR duplicate reads in each group. SnapTools outputs a .snap file, which is an hdf5 file that stores the input scATAC-seq data, including cell-by-bin matrix used in the development and analyses of simATAC modelling [11]. For 10xG samples, we started from the fragment.tsv file provided by 10x website, which is a barcoded and aligned fragment file processed, with an implemented option by SnapTools for 10xG samples. The rest of the samples were processed from FASTQ or provided BAM files, and unique randomly generated barcodes were added to the samples that did not have barcodes themselves. We used samtools (version 1.10) for some of our pre-processing [40]. Bedtools (version 2.27.1) was used for generating peak-by-cell matrices [41], and Picard tool (version 2.23.3) for removing duplicate reads [42]. SnapATAC (version 1.0.0) R package was used for data pre-processing, loading bin-by-cell matrices, and clustering analysis [39]. We also used Signac R package (version 1.0.0), which is an extension of Seurat for the clustering analysis of peak-by-cell matrices [43].

The code and dataset files used for benchmarking are available at https://github.com/bowang-lab/simATAC [44].

## Availability of data and materials

The datasets analyzed during the current study are available from the repositories specified in Methods. The simATAC R package and code used to analyze the data is available under a GPLv3 license on the GitHub repository for this paper https://github.com/bowang-lab/simATAC(doi: 10.5281/zenodo.4411995) [45]. Copies of the benchmark datasets are also provided in this repository.

## Author contributions

ZN developed the software and performed the analyses. LZ contributed to the statistical analyses and writing the manuscript. BW oversaw all aspects of the project. All authors contributed to drafting the manuscript. All authors read and approved the final manuscript.

## Acknowledgements

We would like to thank Chloe Wang, Hassaan Maan, Dr. Mehran Karimzadeh, Ronald Xie, and Rex Ma for their helpful comments on the manuscript.

## Consent for publication

Not applicable.

## Ethics approval and consent to participate

Not applicable.

## Additional Files

Additional file 1 — Figure S1-S15, Table S1-S9, Note S1-S2

Figures of parameter comparison for cell groups that are not provided in the main manuscript, comparison of real and simulated parameters’ polynomial relation for 12 sample cell groups, parameter plots for simATAC simulated data with different parameters’ adjustment, cell type clustering results, table of benchmark datasets’ simulation time, table of correlations, and table of all datasets’ detailed information, table of abbreviations, description of the pipeline used for generating peak-by-cell matrix, and description of clustering parameter equations (provided in PDF format).

Additional file 2

Table of estimated values for Gaussian mixture model and polynomial function parameters obtained for benchmark cell groups (provided in XLSX format).

Additional file 3

Table of statistical analysis parameters of simATAC library size modelling (provided in XLSX format).

## References

1. Olsen, T.K., Baryawno, N.: Introduction to Single-Cell RNA Sequencing. Current Protocols in Molecular Biology (2018). doi:10.1002/cpmb.57

2. Thurman, R.E., Rynes, E., Humbert, R., Vierstra, J., Maurano, M.T., Haugen, E., Sheffield, N.C., Stergachis, A.B., Wang, H., Vernot, B., Garg, K., John, S., Sandstrom, R., Bates, D., Boatman, L., Canfield, T.K., Diegel, M., Dunn, D., Ebersol, A.K., Frum, T., Giste, E., Johnson, A.K., Johnson, E.M., Kutyavin, T., Lajoie, B., Lee, B.K., Lee, K., London, D., Lotakis, D., Neph, S., Neri, F., Nguyen, E.D., Qu, H., Reynolds, A.P., Roach, V., Safi, A., Sanchez, M.E., Sanyal, A., Shafer, A., Simon, J.M., Song, L., Vong, S., Weaver, M., Yan, Y., Zhang, Z., Zhang, Z., Lenhard, B., Tewari, M., Dorschner, M.O., Hansen, R.S., Navas, P.A., Stamatoyannopoulos, G., Iyer, V.R., Lieb, J.D., Sunyaev, S.R., Akey, J.M., Sabo, P.J., Kaul, R., Furey, T.S., Dekker, J., Crawford, G.E., Stamatoyannopoulos, J.A.: The accessible chromatin landscape of the human genome. Nature (2012). doi:10.1038/nature11232

3. Stergachis, A.B., Neph, S., Reynolds, A., Humbert, R., Miller, B., Paige, S.L., Vernot, B., Cheng, J.B., Thurman, R.E., Sandstrom, R., Haugen, E., Heimfeld, S., Murry, C.E., Akey, J.M., Stamatoyannopoulos, J.A.: Developmental fate and cellular maturity encoded in human regulatory DNA landscapes. Cell (2013). doi:10.1016/j.cell.2013.07.020

4. Buenrostro, J.D., Wu, B., Chang, H.Y., Greenleaf, W.J.: ATAC-seq: A method for assaying chromatin accessibility genome-wide. Current Protocols in Molecular Biology (2015). doi:10.1002/0471142727.mb2129s109

5. Jin, S., Zhang, L., Nie, Q.: scAI: an unsupervised approach for the integrative analysis of parallel single-cell transcriptomic and epigenomic profiles. Genome biology (2020). doi:10.1186/s13059-020-1932-8

6. Stuart, T., Butler, A., Hoffman, P., Hafemeister, C., Papalexi, E., Mauck, W.M., Hao, Y., Stoeckius, M., Smibert, P., Satija, R.: Comprehensive Integration of Single-Cell Data. Cell (2019). doi:10.1016/j.cell.2019.05.031

7. Butler, A., Hoffman, P., Smibert, P., Papalexi, E., Satija, R.: Integrating single-cell transcriptomic data across different conditions, technologies, and species. Nature Biotechnology (2018). doi:10.1038/nbt.4096

8. Schep, A.N., Wu, B., Buenrostro, J.D., Greenleaf, W.J.: ChromVAR: Inferring transcription-factor-associated accessibility from single-cell epigenomic data. Nature Methods (2017). doi:10.1038/nmeth.4401

9. Ji, Z., Zhou, W., Ji, H.: Single-cell regulome data analysis by SCRAT. Bioinformatics (2017). doi:10.1093/bioinformatics/btx315

10. Zamanighomi, M., Lin, Z., Daley, T., Chen, X., Duren, Z., Schep, A., Greenleaf, W.J., Wong, W.H.: Unsupervised clustering and epigenetic classification of single cells. Nature Communications (2018). doi:10.1038/s41467-018-04629-3

11. Fang, R., Preissl, S., Li, Y., Hou, X., Lucero, J., Wang, X., Motamedi, A., Shiau, A.K., Zhou, X., Xie, F., Mukamel, E.A., Zhang, K., Zhang, Y., Behrens, M.M., Ecker, J.R., Ren, B.: SnapATAC: A comprehensive analysis package for single cell ATAC-seq. bioRxiv (2020). doi:10.1101/615179

12. Bravo González-Blas, C., Minnoye, L., Papasokrati, D., Aibar, S., Hulselmans, G., Christiaens, V., Davie, K., Wouters, J., Aerts, S.: cisTopic: cis-regulatory topic modeling on single-cell ATAC-seq data. Nature Methods (2019). doi:10.1038/s41592-019-0367-1

13. Urrutia, E., Chen, L., Zhou, H., Jiang, Y.: Destin: toolkit for single-cell analysis of chromatin accessibility. Bioinformatics 35(19), 3818–3820 (2019). doi:10.1093/bioinformatics/btz141

14. Li, B., Li, Y., Li, K., Zhu, L., Yu, Q., Cai, P., Fang, J., Zhang, W., Du, P., Jiang, C., Lin, J., Qu, K.: APEC: an accesson-based method for single-cell chromatin accessibility analysis. Genome biology (2020). doi:10.1186/s13059-020-02034-y

15. Zhao, C., Hu, S., Huo, X., Zhang, Y.: Dr.seq2: A quality control and analysis pipeline for parallel single cell transcriptome and epigenome data. PLoS ONE (2017). doi:10.1371/journal.pone.0180583

16. Xiong, L., Xu, K., Tian, K., Shao, Y., Tang, L., Gao, G., Zhang, M., Jiang, T., Zhang, Q.C.: SCALE method for single-cell ATAC-seq analysis via latent feature extraction. Nature Communications (2019). doi:10.1038/s41467-019-12630-7

17. Chen, H., Lareau, C., Andreani, T., Vinyard, M.E., Garcia, S.P., Clement, K., Andrade-Navarro, M.A., Buenrostro, J.D., Pinello, L.: Assessment of computational methods for the analysis of single-cell ATAC-seq data. Genome Biology 20(1), 241 (2019). doi:10.1186/s13059-019-1854-5

18. Duren, Z., Chen, X., Zamanighomi, M., Zeng, W., Satpathy, A.T., Chang, H.Y., Wang, Y., Wong, W.H.: Integrative analysis of single-cell genomics data by coupled nonnegative matrix factorizations. Proceedings of the National Academy of Sciences of the United States of America (2018). doi:10.1073/pnas.1805681115

19. de Boer, C.G., Regev, A.: BROCKMAN: Deciphering variance in epigenomic regulators by k-mer factorization. BMC Bioinformatics (2018). doi:10.1186/s12859-018-2255-6

20. Li, Z., Schulz, M.H., Look, T., Begemann, M., Zenke, M., Costa, I.G.: Identification of transcription factor binding sites using ATAC-seq. Genome Biology (2019). doi:10.1186/s13059-019-1642-2

21. Lun, A., Risso, D.: SingleCellExperiment: S4 Classes for Single Cell Data. (2020). R package version 1.10.1

22. Buenrostro, J.D., Corces, M.R., Lareau, C.A., Wu, B., Schep, A.N., Aryee, M.J., Majeti, R., Chang, H.Y., Greenleaf, W.J.: Integrated Single-Cell Analysis Maps the Continuous Regulatory Landscape of Human Hematopoietic Differentiation. Cell (2018). doi:10.1016/j.cell.2018.03.074

23. Cusanovich, D.A., Hill, A.J., Aghamirzaie, D., Daza, R.M., Pliner, H.A., Berletch, J.B., Filippova, G.N., Huang, X., Christiansen, L., DeWitt, W.S., Lee, C., Regalado, S.G., Read, D.F., Steemers, F.J., Disteche, C.M., Trapnell, C., Shendure, J.: A Single-Cell Atlas of In Vivo Mammalian Chromatin Accessibility. Cell (2018). doi:10.1016/j.cell.2018.06.052

24. 10x Genomics. 5k Peripheral blood mononuclear cells (PBMCs) from a healthy donor. Accessed: 2021-01-08. https://support.10xgenomics.com/single-cell-atac/datasets/1.0.1/atac_v1_pbmc_5k

25. Zhang, Y., Liu, T., Meyer, C.A., Eeckhoute, J., Johnson, D.S., Bernstein, B.E., Nusbaum, C., Myers, R.M., Brown, M., Li, W., Liu, X.S.: Model-based Analysis of ChIP-Seq (MACS). Genome Biology 9(9), 137 (2008). doi:10.1186/gb-2008-9-9-r137

26. Gaspar, J.M.: Genrich. GitHub. Accessed: 2021-01-08 (2019). https://github.com/jsh58/Genrich

27. R Core Team: R: A Language and Environment for Statistical Computing. R Foundation for Statistical Computing, Vienna, Austria (2019). R Foundation for Statistical Computing. https://www.R-project.org/

28. Delignette-Muller, M.L., Dutang, C.: fitdistrplus: An R package for fitting distributions. Journal of Statistical Software 64(4), 1–34 (2015)

29. Maechler, M.: Diptest: Hartigan’s Dip Test Statistic for Unimodality - Corrected. (2016). R package version 0.75-7. https://CRAN.R-project.org/package=diptest

30. Benaglia, T., Chauveau, D., Hunter, D.R., Young, D.: mixtools: An R package for analyzing finite mixture models. Journal of Statistical Software 32(6), 1–29 (2009)

31. Pedregosa, F., Varoquaux, G., Gramfort, A., Michel, V., Thirion, B., Grisel, O., Blondel, M., Prettenhofer, P., Weiss, R., Dubourg, V., Vanderplas, J., Passos, A., Cournapeau, D., Brucher, M., Perrot, M., Duchesnay, E.: Scikit-learn: Machine learning in Python. Journal of Machine Learning Research 12, 2825–2830 (2011)

32. Satpathy, A.T., Granja, J.M., Yost, K.E., Qi, Y., Meschi, F., McDermott, G.P., Olsen, B.N., Mumbach, M.R., Pierce, S.E., Corces, M.R., Shah, P., Bell, J.C., Jhutty, D., Nemec, C.M., Wang, J., Wang, L., Yin, Y., Giresi, P.G., Chang, A.L.S., Zheng, G.X.Y., Greenleaf, W.J., Chang, H.Y.: Massively parallel single-cell chromatin landscapes of human immune cell development and intratumoral T cell exhaustion. Nature Biotechnology 37(8), 925–936 (2019). doi:10.1038/s41587-019-0206-z

33. Corces, M.R., Buenrostro, J.D., Wu, B., Greenside, P.G., Chan, S.M., Koenig, J.L., Snyder, M.P., Pritchard, J.K., Kundaje, A., Greenleaf, W.J., Majeti, R., Chang, H.Y.: Lineage-specific and single-cell chromatin accessibility charts human hematopoiesis and leukemia evolution. Nature Genetics (2016). doi:10.1038/ng.3646

34. Buenrostro, J.D., Wu, B., Litzenburger, U.M., Ruff, D., Gonzales, M.L., Snyder, M.P., Chang, H.Y., Greenleaf, W.J.: Single-cell chromatin accessibility reveals principles of regulatory variation. Nature (2015). doi:10.1038/nature14590

35. Cusanovich, D.A., Daza, R., Adey, A., Pliner, H.A., Christiansen, L., Gunderson, K.L., Steemers, F.J., Trapnell, C., Shendure, J.: Multiplex single-cell profiling of chromatin accessibility by combinatorial cellular indexing. Science (2015). doi:10.1126/science.aab1601

36. Chen, X., Litzenburger, U.M., Wei, Y., Schep, A.N., LaGory, E.L., Choudhry, H., Giaccia, A.J., Greenleaf, W.J., Chang, H.Y.: Joint single-cell DNA accessibility and protein epitope profiling reveals environmental regulation of epigenomic heterogeneity. Nature Communications (2018). doi:10.1038/s41467-018-07115-y

37. Preissl, S., Fang, R., Huang, H., Zhao, Y., Raviram, R., Gorkin, D.U., Zhang, Y., Sos, B.C., Afzal, V., Dickel, D.E., Kuan, S., Visel, A., Pennacchio, L.A., Zhang, K., Ren, B.: Single-nucleus analysis of accessible chromatin in developing mouse forebrain reveals cell-type-specific transcriptional regulation. Nature Neuroscience (2018). doi:10.1038/s41593-018-0079-3

38. 10x Genomics. Cell Ranger ATAC. Accessed: 2021-01-08. https://support.10xgenomics.com/single-cell-atac/software/pipelines/latest/what-is-cell-ranger-atac

39. Fang, R.: SnapATAC: Single Nucleus Analysis Package for ATAC-Seq. (2019). R package version 1.0.0. https://github.com/r3fang/SnapATAC

40. Li, H., Handsaker, B., Wysoker, A., Fennell, T., Ruan, J., Homer, N., Marth, G., Abecasis, G., Durbin, R.: The sequence alignment/map format and samtools. Bioinformatics 25(16), 2078–2079 (2009)

41. Quinlan, A.R., Hall, I.M.: Bedtools: a flexible suite of utilities for comparing genomic features. Bioinformatics 26(6), 841–842 (2010)

42. Picard toolkit. Broad Institute. Accessed: 2021-01-08 (2019). http://broadinstitute.github.io/picard/

43. Stuart, T., Srivastava, A., Lareau, C., Satija, R.: Multimodal single-cell chromatin analysis with signac. bioRxiv (2020). doi:10.1101/2020.11.09.373613

44. Navidi, Z., Zhang, L., Wang, B.: simATAC: a single-cell ATAC-seq simulation framework. GitHub (2020). https://github.com/bowang-lab/simATAC

45. Navidi, Z., Zhang, L., Wang, B.: simATAC: a single-cell ATAC-seq simulation framework. Zenodo (2020). https://doi.org/10.5281/zenodo.4411995

