## Additional file 1 for "simATAC: a single-cell ATAC-seq simulation framework"

### simATAC Additional File 1

##### ADDITIONAL FIGURES, TABLES, AND NOTES

**Figure S1.** Library size box plots

**Figure S2.** Bin sparsity QQ-plots

**Figure S3.** Cell sparsity box plots

**Figure S4.** Bin sparsity QQ-plots for sparsity adjustment factor of 1

**Figure S5.** Bin sparsity QQ-plots for sparsity adjustment factor of 0.9

**Figure S6.** Bin sparsity QQ-plots for sparsity adjustment factor of 0.8

**Figure S7.** Parameters' demonstration for different sparsity adjustment factors

**Figure S8.** Peak-by-cell clustering results

**Figure S9.** Buenrostro2018 plots (Noise mean: -0.3, noise sd: 0.3)

**Figure S10.** Buenrostro2018 plots (Noise mean: -0.4, noise sd: 0.4)

**Figure S11.** Cusanovich2018 plots (Noise mean: -0.3, noise sd: 0.3)

**Figure S12.** Cusanovich2018 plots (Noise mean: -0.4, noise sd: 0.4)

**Figure S13.** PBMCs plots (Noise mean: -0.3, noise sd: 0.3)

**Figure S14.** PBMCs plots (Noise mean: -0.4, noise sd: 0.4)

**Figure S15.** Polynomial regression relationship between bin means and non-zero cell proportions

**Table S1.** Table of real scATAC-seq datasets

**Table S2.** Table of correlation (No noise)

**Table S3.** Table of peak calling comparison

**Table S4.** Table of peak-by-cell clustering comparison

**Table S5.** Table of correlation (Noise mean: -0.3, noise sd: 0.3)

**Table S6.** Table of correlation (Noise mean: -0.4, noise sd: 0.4)

**Table S7.** Table of clustering results for different sparsity adjustment factor

**Table S8.** Table of simATAC running time

**Table S9.** Table of abbreviations

**Note S1.** Peak-by-cell matrix generation pipeline

**Note S2.** Clustering metrics calculation

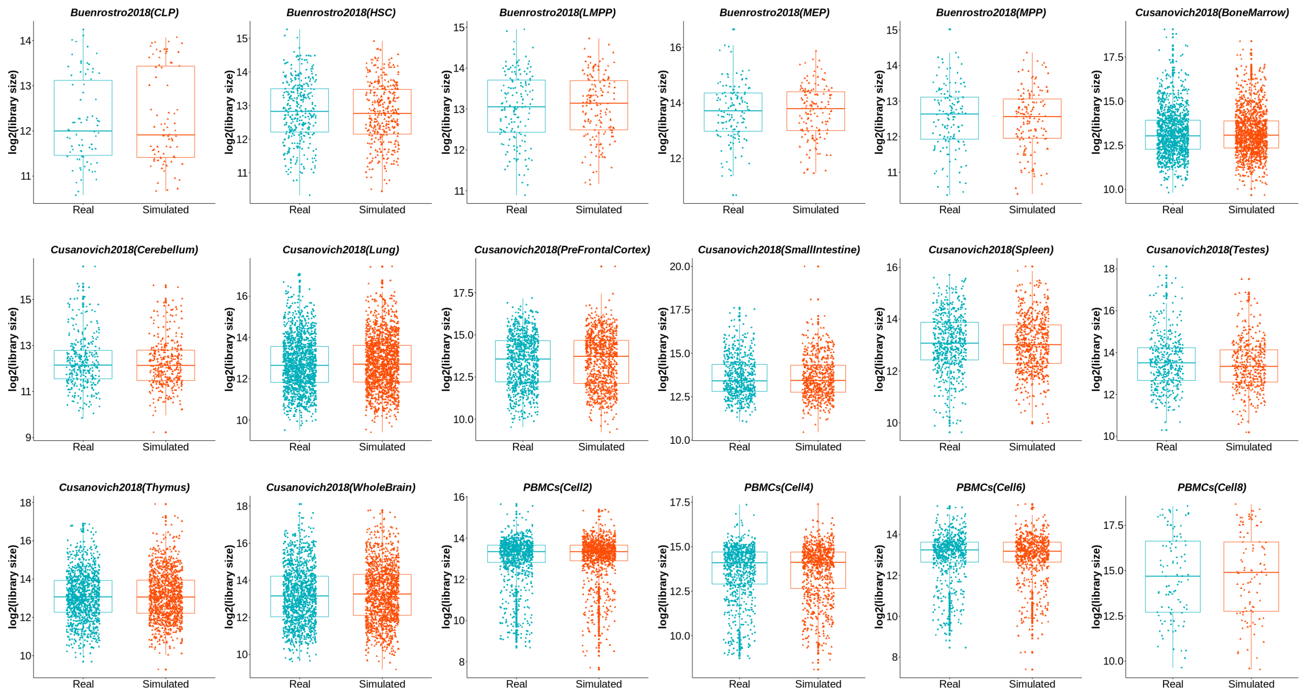


**Figure 1.** Library size box plot demonstration of real and simATAC simulated bin-by-cell matrices for the cell groups from benchmark datasets: Buenrostro2018, Cusanovich2018, and PBMCs.

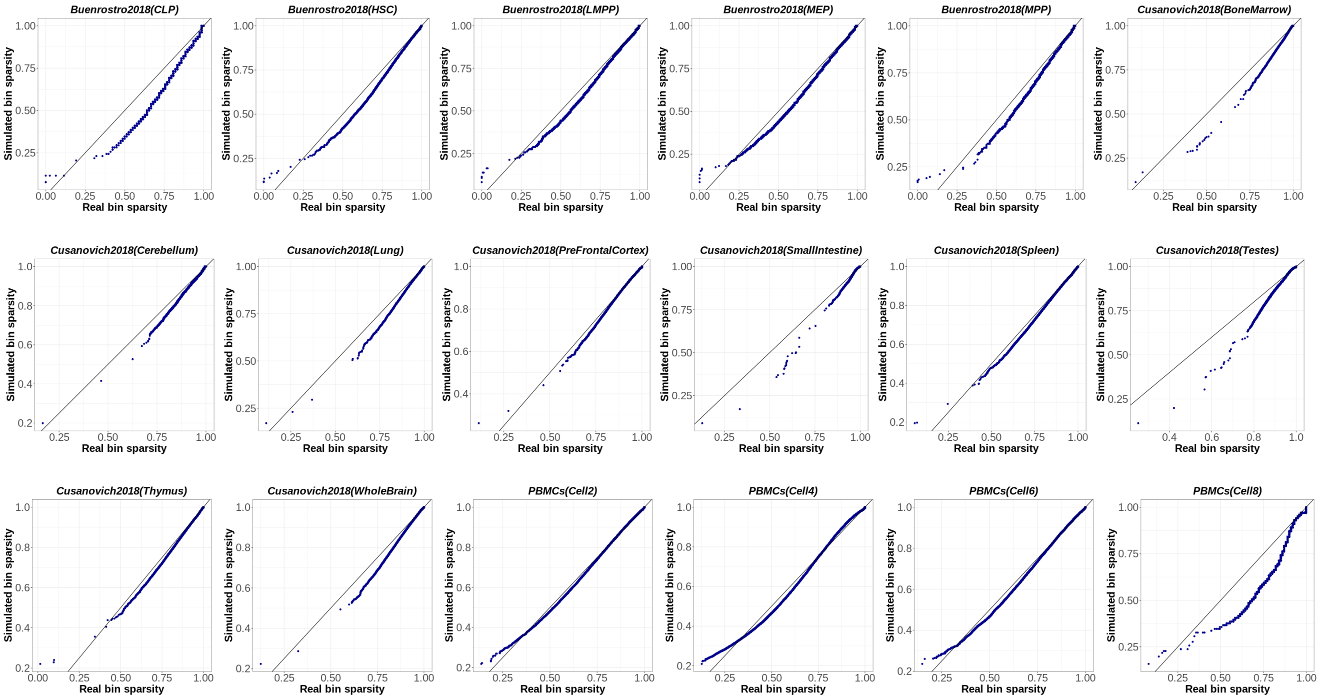


**Figure 2.** Bin sparsity QQ-plot demonstration of real and simATAC simulated bin-by-cell matrices for the cell groups from benchmark datasets: Buenrostro2018, Cusanovich2018, and PBMCs.

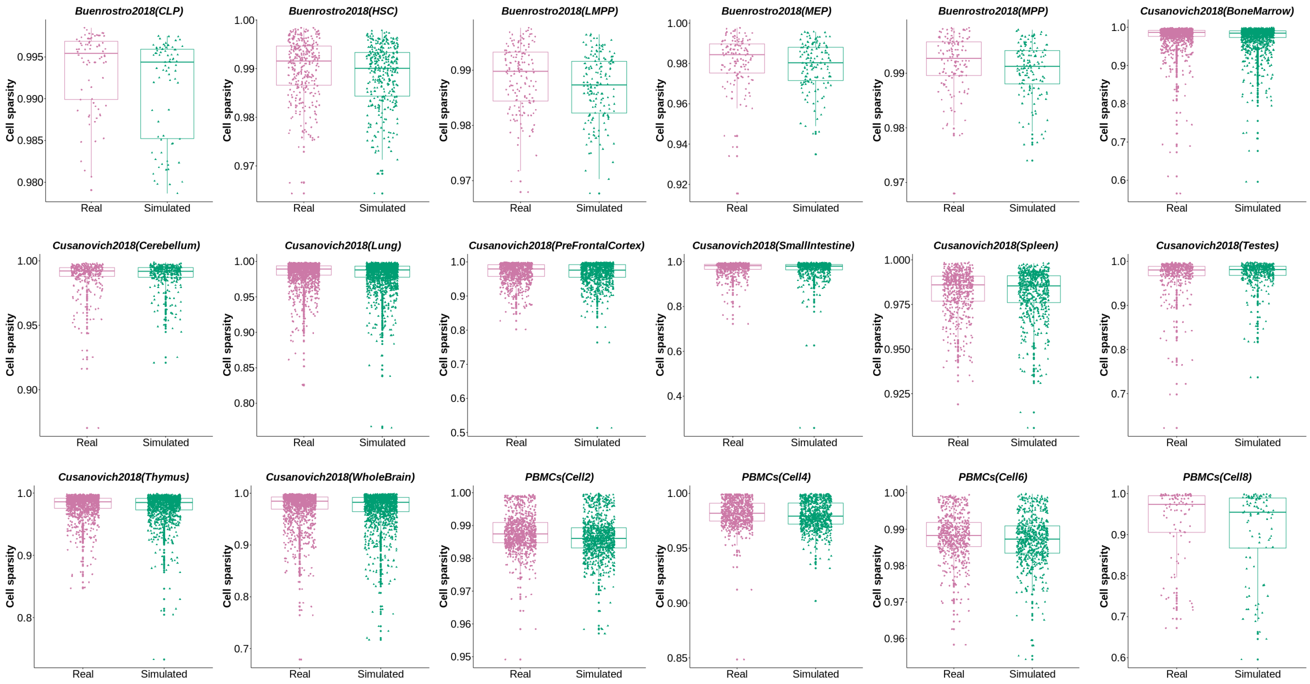


**Figure 3.** Cell sparsity box plot demonstration of real and simATAC simulated bin-by-cell matrices for the cell groups from benchmark datasets: Buenrostro2018, Cusanovich2018, and PBMCs.

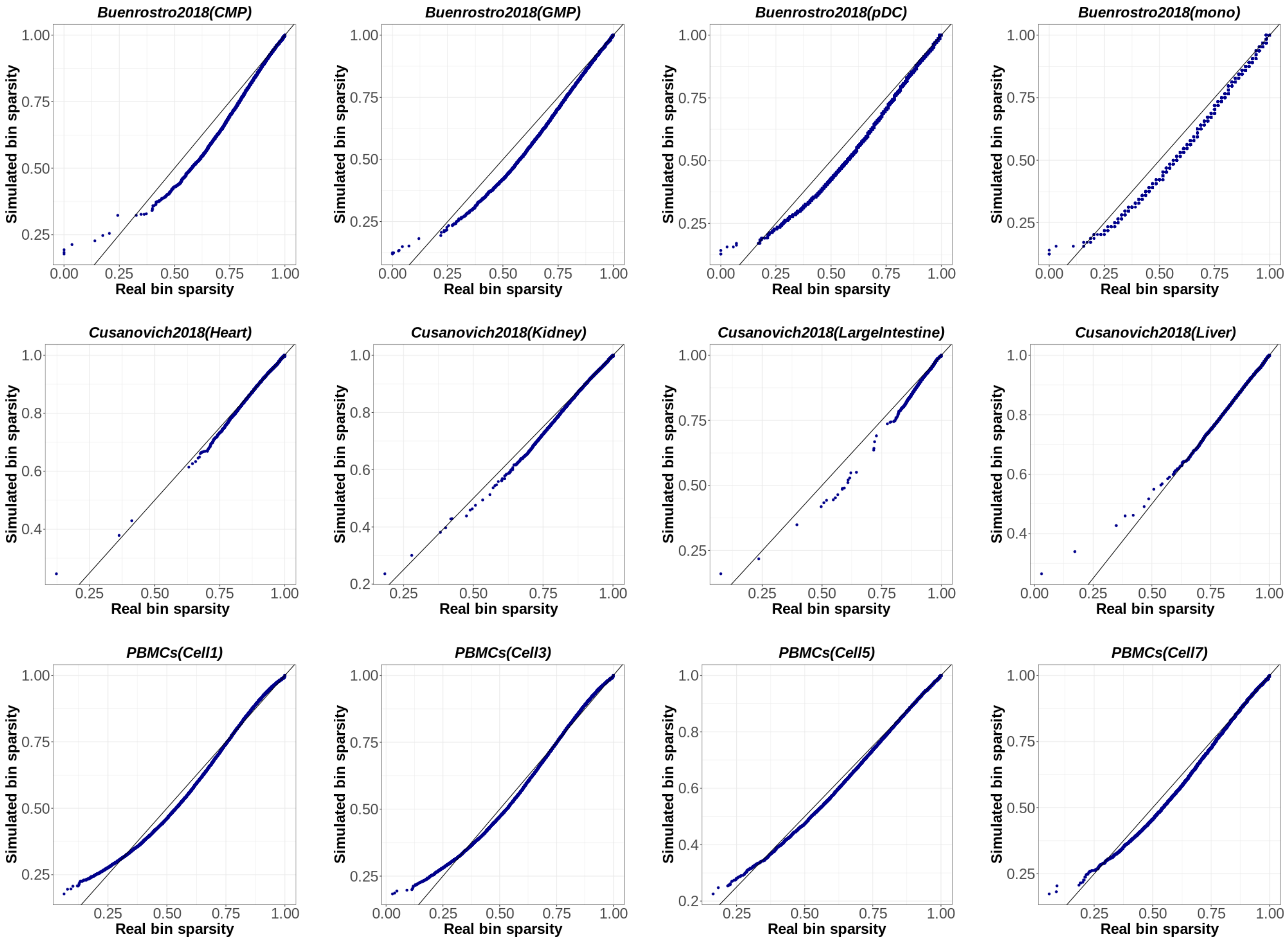


**Figure 4.** Bin sparsity QQ-plots of real and simATAC simulated bin-by-cell matrices with a sparsity adjustment factor of 1, demonstrated for the 12 sample cell groups from benchmark datasets: Buenrostro2018, Cusanovich2018, and PBMCs.

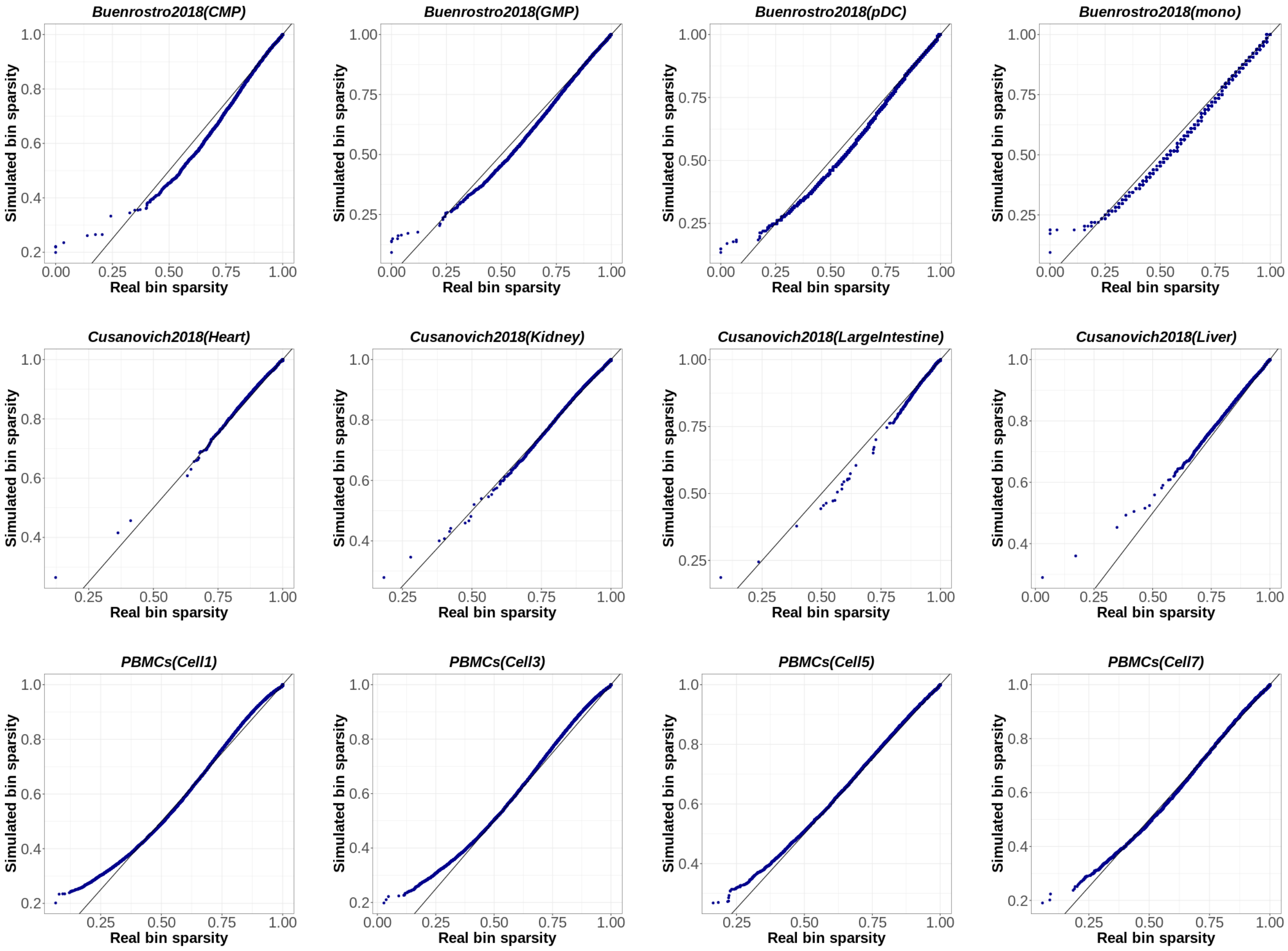


**Figure 5.** Bin sparsity QQ-plots of real and simATAC simulated bin-by-cell matrices with a sparsity adjustment factor of 0.9, demonstrated for the 12 sample cell groups from benchmark datasets: Buenrostro2018, Cusanovich2018, and PBMCs.

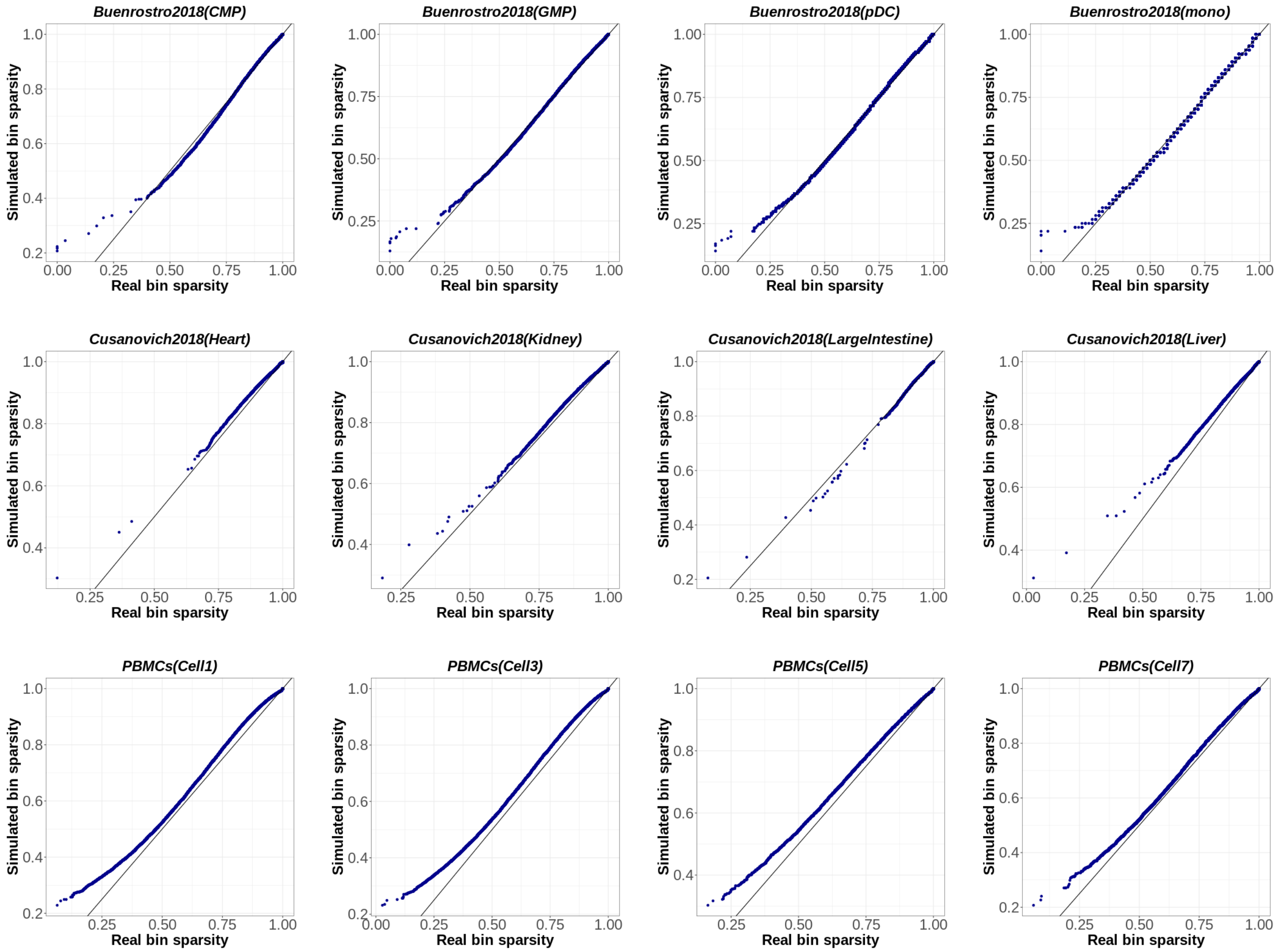


**Figure 6.** Bin sparsity QQ-plots of real and simATAC simulated bin-by-cell matrices with a sparsity adjustment factor of 0.8, demonstrated for the 12 sample cell groups from benchmark datasets: Buenrostro2018, Cusanovich2018, and PBMCs.

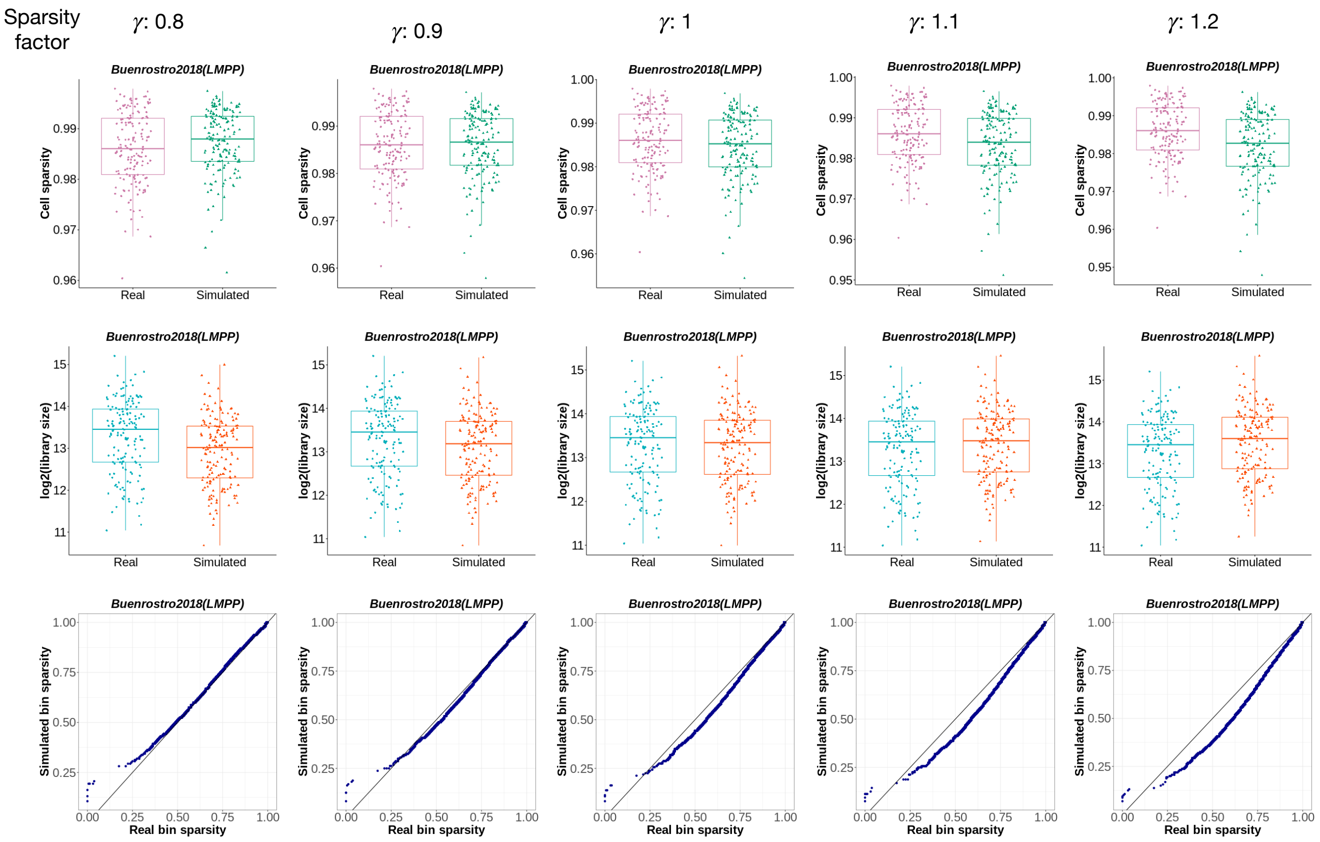


**Figure 7.** Cell sparsity, library size, and bin sparsity demonstration of real and simATAC simulated bin-by-cell matrices for different values of the sparsity adjustment factor $\gamma$ in $\{0.8, 0.9, 1, 1.1, 1.2\}$ for the Buenrostro2018 dataset LMPP cell type.

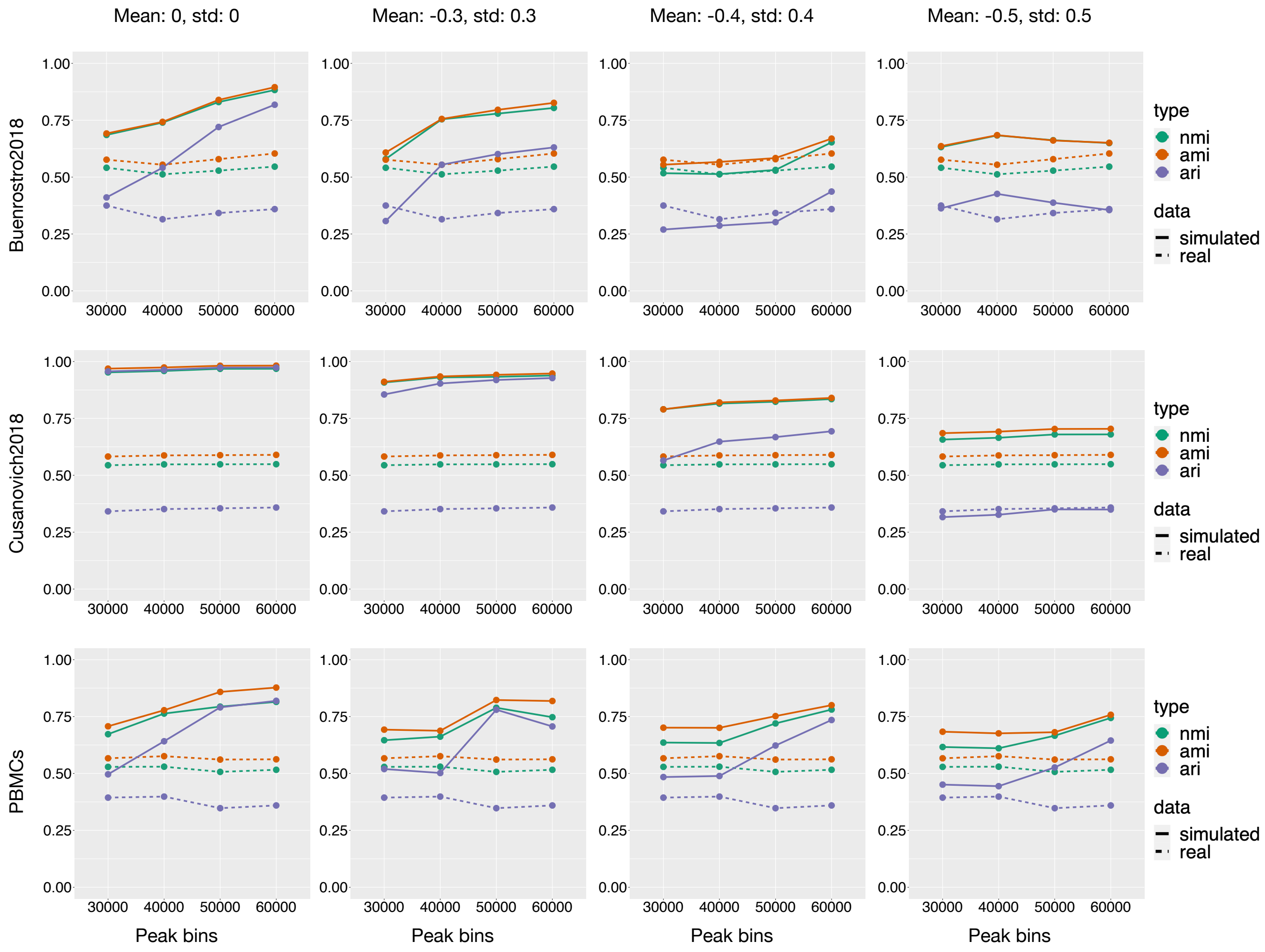


**Figure 8.** Clustering performance comparison of peak-by-cell matrices extracted from the real and simulated bin-by-cell arrays. Twenty simulation runs were performed on each benchmark dataset considering different Gaussian noise levels (mean and standard deviation) and different numbers of peak bins. Averaged NMI, AMI, and ARI metrics obtained from Seurat's graph-based clustering algorithm are demonstrated for different called peak bin numbers and noise levels.

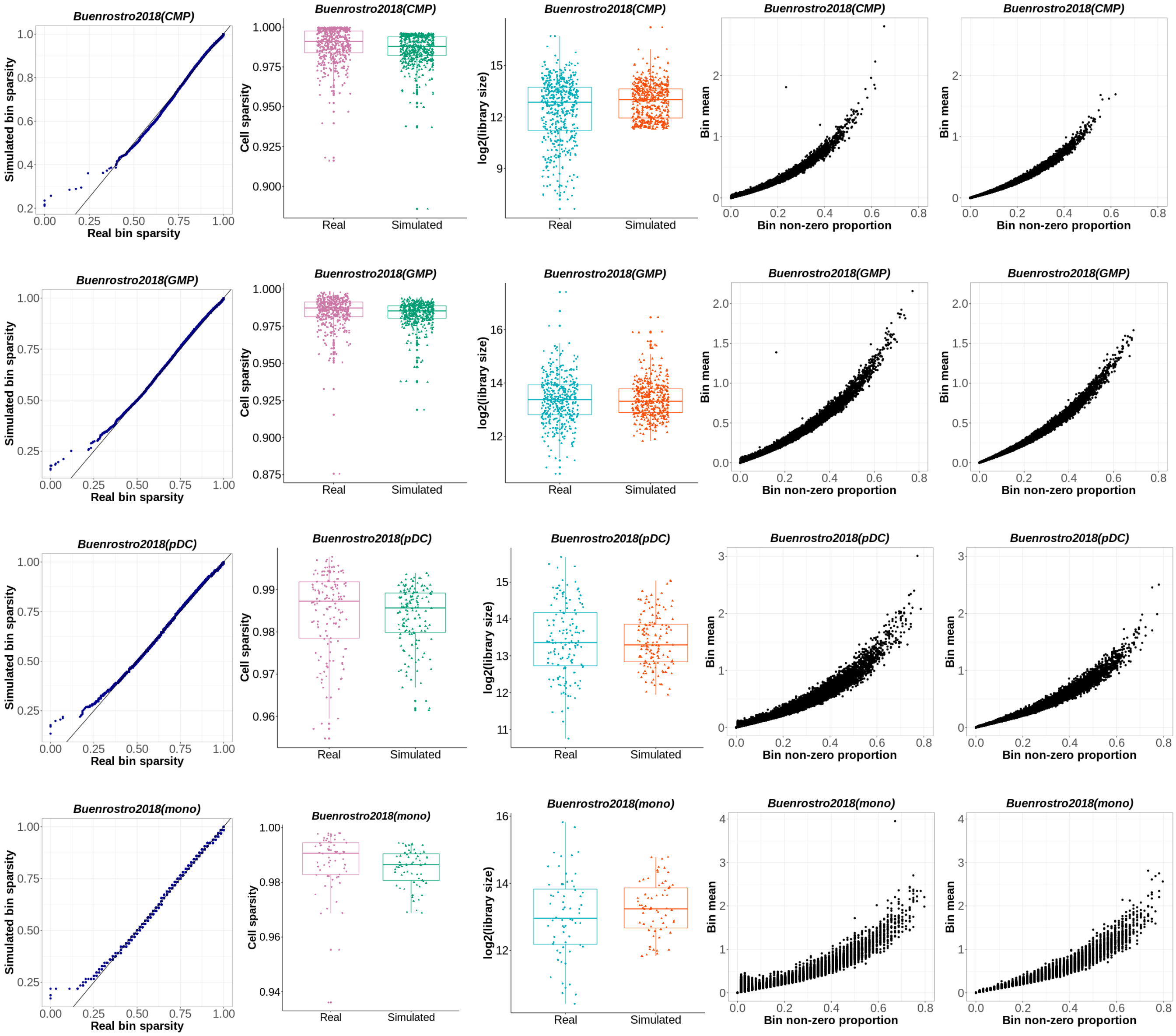


**Figure 9.** Buenrostro2018 parameters' plots for Gaussian noise mean: -0.3, and standard deviation: 0.3. Bin sparsity QQ-plot, cell sparsity box plot, library size box plot, real matrix bin means and non-zero cell proportions relationship (excluding bins with non-zero cell proportion $\geq 0.8$), simulated matrix bin means and non-zero cell proportions relationship (excluding bins with non-zero cell proportion $\geq 0.8$).

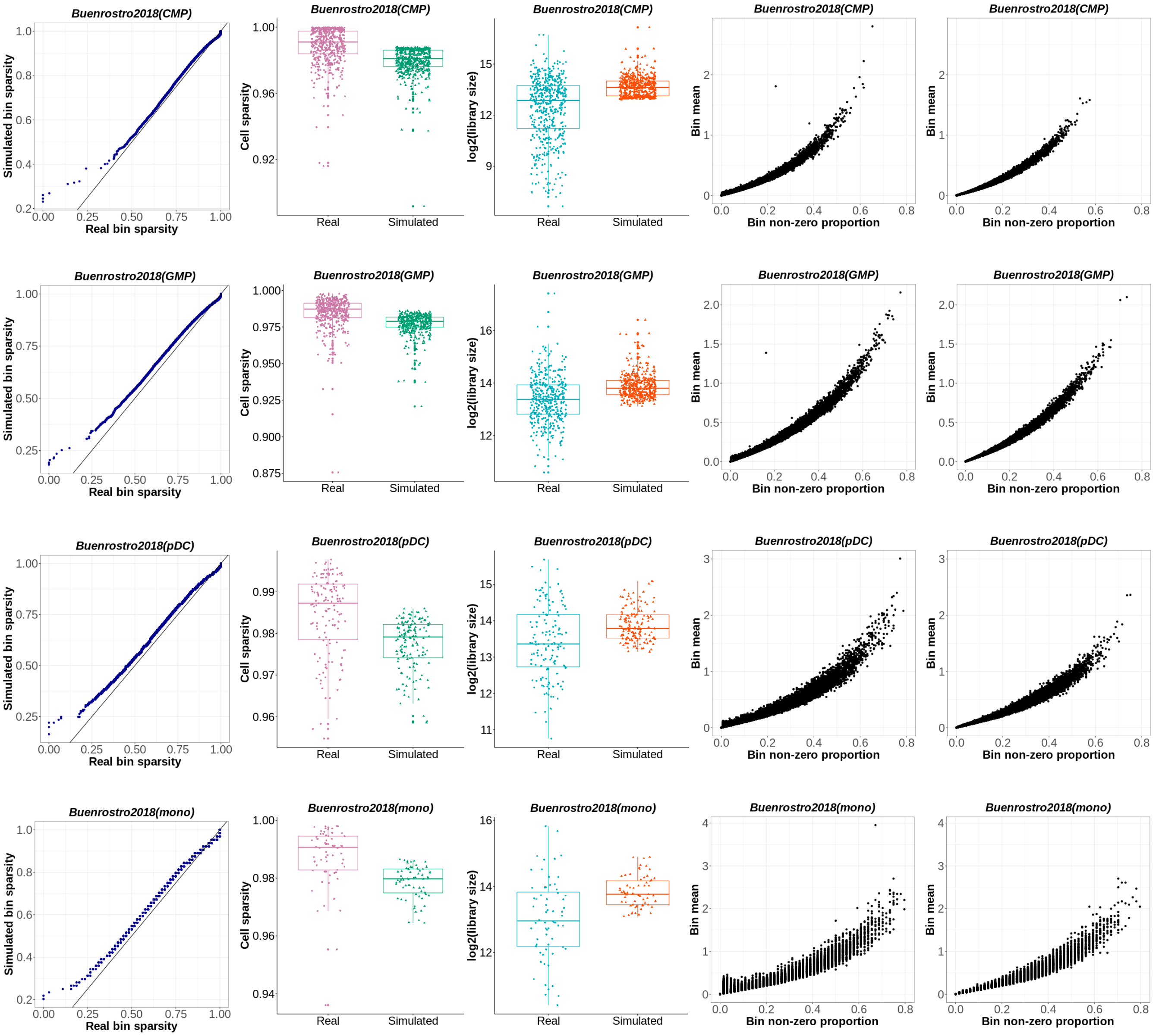


**Figure 10.** Buenrostro2018 parameters' plots for Gaussian noise mean: -0.4, and standard deviation: 0.4. Bin sparsity QQ-plot, cell sparsity box plot, library size box plot, real matrix bin means and non-zero cell proportions relationship (excluding bins with non-zero cell proportion $\geq 0.8$), simulated matrix bin means and non-zero cell proportions relationship (excluding bins with non-zero cell proportion $\geq 0.8$).

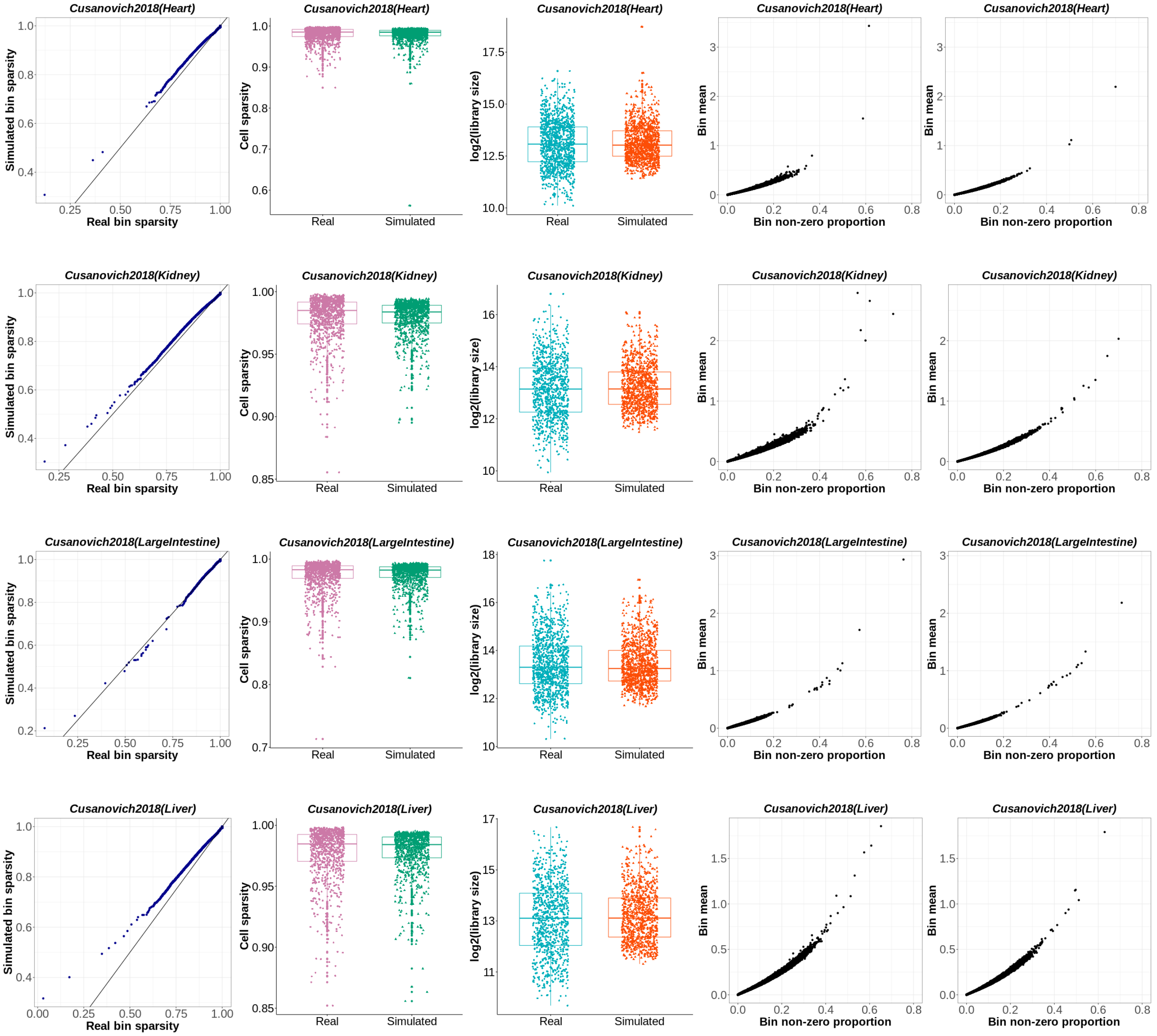


**Figure 11.** Cusanovich2018 parameters' plots for Gaussian noise mean: -0.3, and standard deviation: 0.3. Bin sparsity QQ-plot, cell sparsity box plot, library size box plot, real matrix bin means and non-zero cell proportions relationship (excluding bins with non-zero cell proportion $\geq 0.8$), simulated matrix bin means and non-zero cell proportions relationship (excluding bins with non-zero cell proportion $\geq 0.8$).

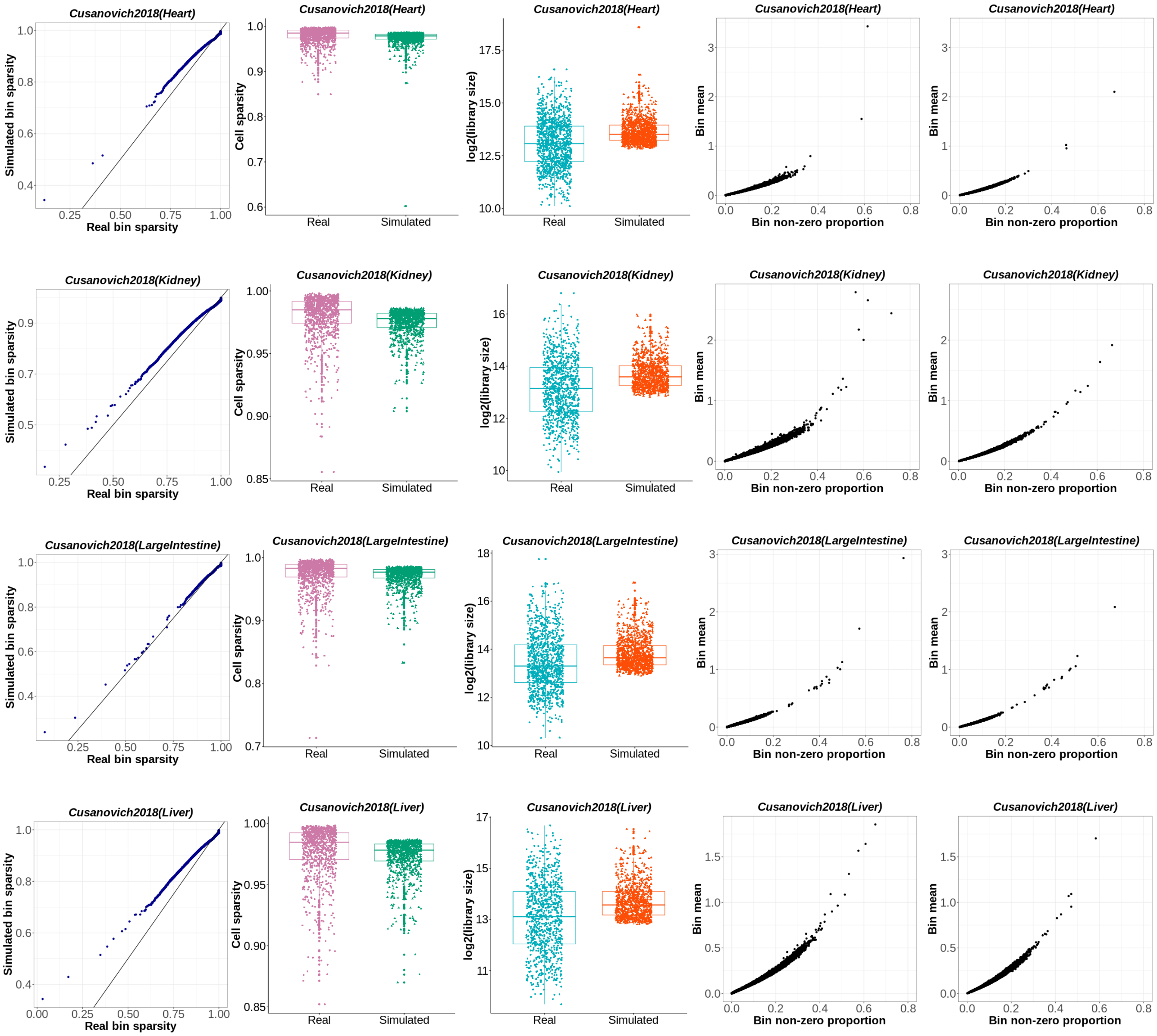


**Figure 12.** Cusanovich2018 parameters' plots for Gaussian noise mean: -0.4, and standard deviation: 0.4. Bin sparsity QQ-plot, cell sparsity box plot, library size box plot, real matrix bin means and non-zero cell proportions relationship (excluding bins with non-zero cell proportion $\geq 0.8$), simulated matrix bin means and non-zero cell proportions relationship (excluding bins with non-zero cell proportion $\geq 0.8$).

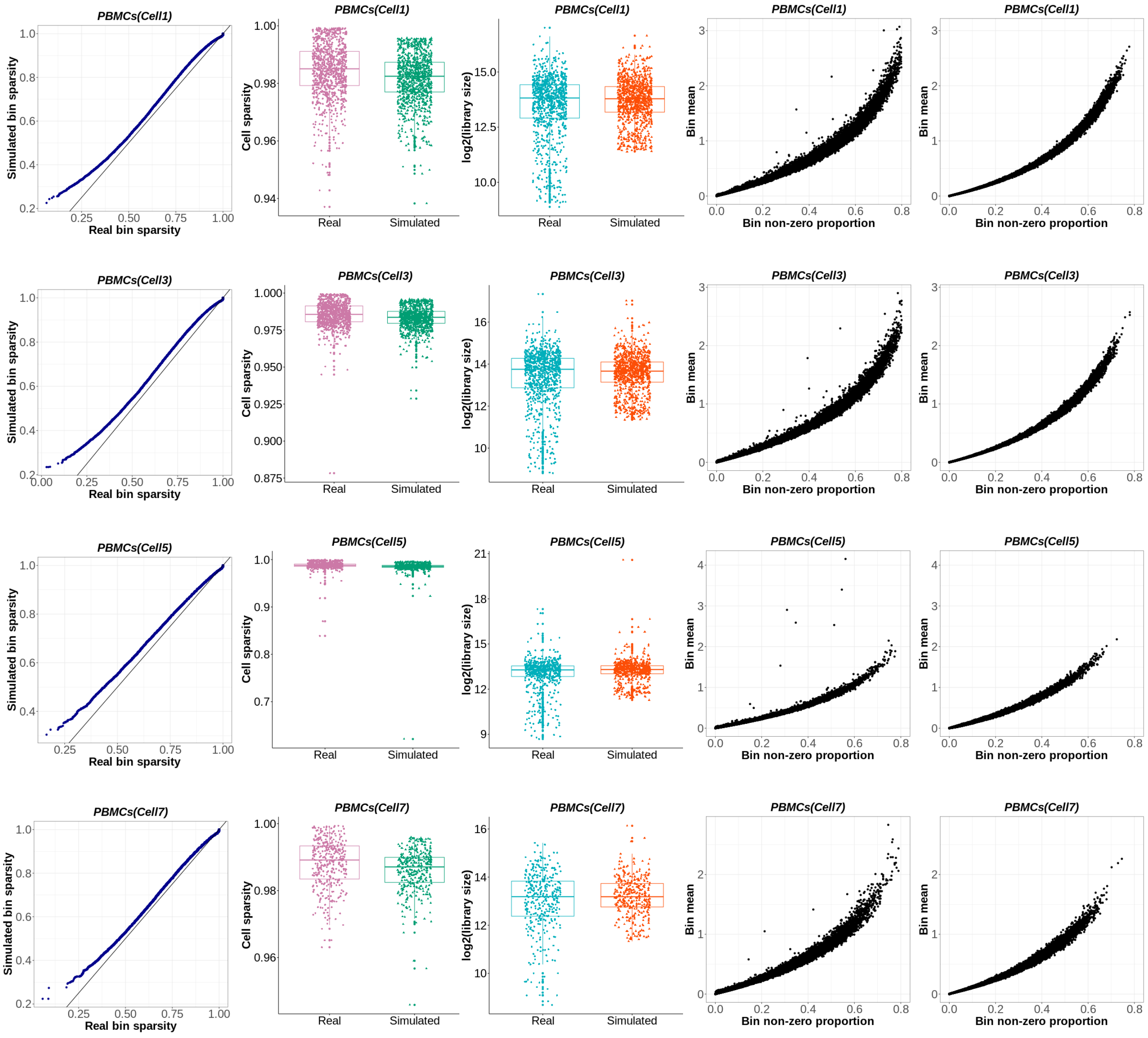


**Figure 13.** PBMCs parameters' plots for Gaussian noise mean: -0.3, and standard deviation: 0.3. Bin sparsity QQ-plot, cell sparsity box plot, library size box plot, real matrix bin means and non-zero cell proportions relationship (excluding bins with non-zero cell proportion $\geq 0.8$), simulated matrix bin means and non-zero cell proportions relationship (excluding bins with non-zero cell proportion $\geq 0.8$).

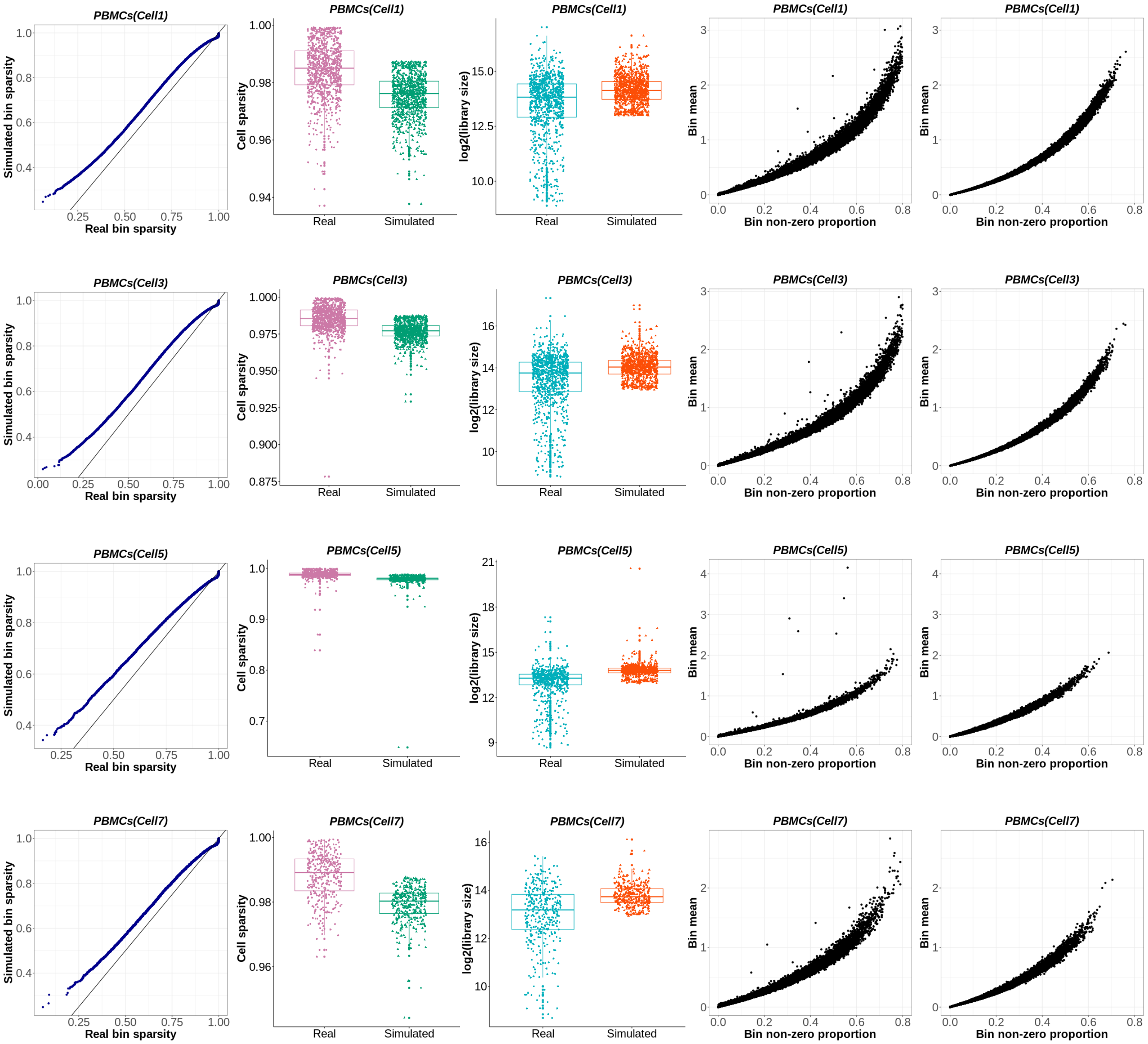


**Figure 14.** PBMCs parameters' plots for Gaussian noise mean: -0.4, and standard deviation: 0.4. Bin sparsity QQ-plot, cell sparsity box plot, library size box plot, real matrix bin means and non-zero cell proportions relationship (excluding bins with non-zero cell proportion $\geq 0.8$), simulated matrix bin means and non-zero cell proportions relationship (excluding bins with non-zero cell proportion $\geq 0.8$).

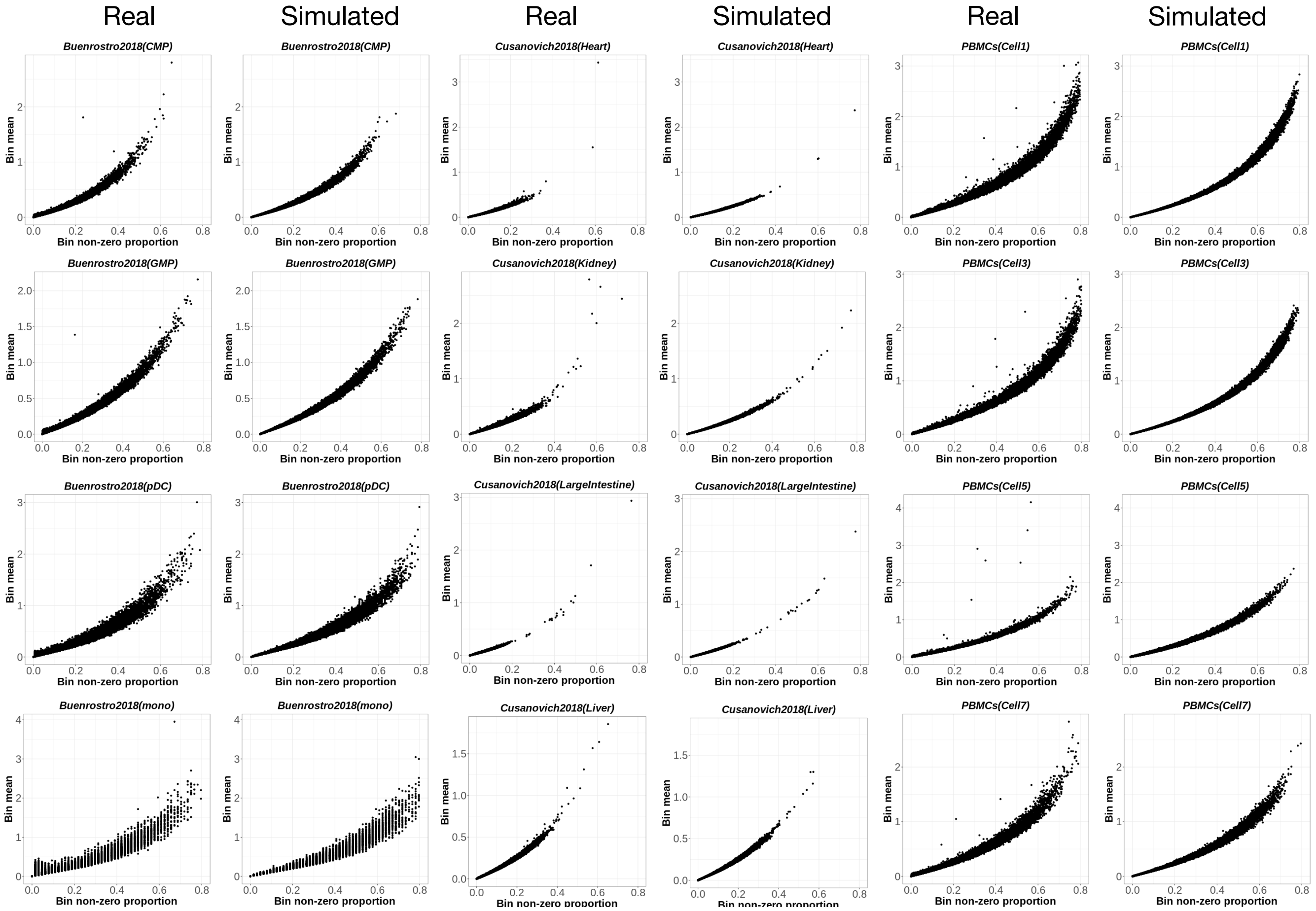


**Figure 15.** The relation between bin means and non-zero cell proportions in real and synthesized count matrices generated by simATAC, demonstrated for the 12 sample cell groups from Buenrostro2018, Cusanovich2018, and PBMCs datasets. The polynomial regression function modelled from real data is used to generate synthetic counts. Bins having a non-zero cell proportion $\geq 0.8$ in the real input data are not included in the plots. Equation 4 in the manuscript generally represents the relationship between bin means $m'_j$ and non-zero cell proportions $p'_j$ when $p'_j < 0.8$. When $p'_j \geq 0.8$, we observed in the modelling real scATAC-seq datasets that equation 4 does not capture this relationship. However, such regions only consist of on average two bins out of $\sim$600,000 bins, and simATAC framework excludes such bins in fitting the polynomial regression model.

**Table 1.** Information of datasets used for modelling and assessing the simATAC framework

| Data | Species | Platform | Cell Groups |
|---|---|---|---|
| GSE99172[1] | Homo sapiens | Fluidigm C1 | Chronic myelogenous leukemia (288) |
| GSE74310[2] | Homo sapiens | Fluidigm C1 | Acute myeloid leukemia-blast cell (192), Acute myeloid leukemia-leukmeia stem cell (191), Lymphoid primed multi-potent progenitor (96), Monocyte (96) |
| GSE65360[3] | Homo sapiens; Mus musculus | Fluidigm C1 | Promyelocytic leukemia cells (96), Human embryonic stem cell line (15), Fibroblasts (96), Lymphoblastoid cells (419), Chronic myeloid leukemia cells (841), Erythroleukemia cell line (96) |
| GSE68103 (GSM1647122)[4] | Homo sapiens | single-cell combinatorial indexing (sciATAC-seq) | HEK293T (232), GM12878 (275), Mixed (19) |
| GSE68103 (GSM1647123)[4] | Homo sapiens | single-cell combinatorial indexing (sciATAC-seq) | GM12878 (334), HL60 (226), Mixed (37) |
| GSE112091 (series GSE112245)[5] | Mus musculus | Multi-Index single-cell ATAC-seq (MI-ATAC) | Epcam+ (192), CD45+ (192) |
| GSE100033 (GSM2668124)[6] | Mus musculus | Single-nucleus ATAC-seq (snATAC-seq) | IN1 (195), EX3 (519), AC (120), EX2 (366), MG (126), EX1 (190), OC (252), IN2 (320) |

| | | | |
|---|---|---|---|
| GSE129785[7] | Homo sapiens; Mus musculus | 10x Genomics Chromium (10xG) | GSM3722011_0p1_99p9_CD4Mem_CD8Naive.snap (7654), GSM3722012_0p1_99p9_Mono_T.snap (10846), GSM3722013_0p5_99p5_CD4Mem_CD8Naive.snap (23551), GSM3722014_0p5_99p5_Mono_T.snap (52323), GSM3722015_PBMC_Rep1.snap (72494), GSM3722016_1_99_CD4Mem_CD8Naive.snap (23058), GSM3722017_1_99_Mono_T.snap (30211), GSM3722018_50_50_CD4Mem_CD8Naive.snap (3212), GSM3722019_50_50_Mono_T.snap (6277), GSM3722020_99_1_CD4Mem_CD8Naive.snap (76717), GSM3722021_99_1_MonoT.snap (86580), GSM3722022_99p5_0p5_CD4Mem_CD8Naive.snap (76896), GSM3722023_99p5_0p5_MonoT.snap (88176), GSM3722024_99p9_0p1_CD4Mem_CD8Naive.snap (79202), GSM3722025_99p9_0p1_Mono_T.snap (84962), GSM3722026_Dendritic_Cells.snap (19562), GSM3722027_Monocytes.snap (28316), GSM3722028_B_Cells.snap (4523), GSM3722029_CD34_Progenitors_Rep1.snap (48069), GSM3722030_Regulatory_T_Cells.snap (3310), GSM3722031_Naive_CD4_T_Cells_Rep1.snap (1658), GSM3722032_Memory_CD4_T_Cells_Rep1.snap (4456), GSM3722033_CD4_HelperT.snap (2060), GSM3722034_CD4_Memory.snap (2981), GSM3722035_CD4_Naive.snap (3489), GSM3722036_NK_Cells.snap (1720), GSM3722037_Naive_CD8_T_Cells.snap (2469), GSM3722038_Memory_CD8_T_Cells.snap (2984), GSM3722039_Dendritic_all_cells.snap (4472), GSM3722040_Fresh_pbmc_5k.snap (56410), GSM3722041_Frozen_sorted_pbmc_5k.snap (61133), GSM3722042_Frozen_unsorted_pbmc_5k.snap (63957), GSM3722047_SU001_Immune_Post2.snap (67504), GSM3722048_SU001_Tcell_Post2.snap (2942), GSM3722049_SU001_Tcell_Post.snap (16750), GSM3722050_SU001_Total_Post2.snap (59469), GSM3722051_SU001_Total_Pre.snap (65016), GSM3722052_SU001_Tumor_Immune_Post.snap (64744), GSM3722053_SU005_Total_Post.snap (55925), GSM3722054_SU006_Immune_Pre.snap (63337), GSM3722055_SU006_Tcell_Pre.snap (35760), GSM3722056_SU006_Total_Post.snap (42630), GSM3722057_SU006_Tumor_Pre.snap (64776), GSM3722058_SU007_Total_Post.snap (49997), GSM3722059_SU008_Immune_Post.snap (72581), GSM3722060_SU008_Immune_Pre.snap (64712), GSM3722061_SU008_Tcell_Post.snap (64276), GSM3722062_SU008_Tcell_Pre.snap (23958), GSM3722063_SU008_Tumor_Post.snap (9838), GSM3722064_SU008_Tumor_Pre.snap (69002), GSM3722065_SU009_Tcell_Post.snap (67291) |
| 10X PBMCs [1] (Benchmark data) | Homo sapiens | 10x Genomics Chromium (10xG) | Cell1 (963), Cell2 (939), Cell3 (928), Cell4 (789), Cell5 (634), Cell6 (619), Cell7 (362), Cell8 (101) |

[1] https://github.com/pinellolab/scATAC-benchmarking/tree/master/Real_Data/10x_PBMC_5k

| | | | |
|---|---|---|---|
| Buenrostro2018[8] [2] (Benchmark data) | Homo sapiens | Fluidigm C1 | Hematopoietic stem cells (HSCs) (347), Multipotent progenitors (MPPs) (142), Lymphoid-primed multipotent progenitors (LMPPs) (160), Common myeloid progenitors (CMPs) (502), Granulocyte-macrophage progenitors (GMPs) (402), Megakaryocyte-erythrocyte progenitors (MEPs) (138), Common lymphoid progenitors (CLPs) (78), Plasmacytoid dendritic cells (pDCs) (141), and Monocytes (mono) (64) |
| Cusanovich2018 subset[9, 10] [3] (Benchmark data) | Mus musculus | single-cell combinatorial indexing (sciATAC-seq) | Bone marrow (1261), Cerebellum (342), Heart (1148), Kidney (965), Large intestine (1063), Liver (925), Lung (1499), Prefrontal cortex (894), Small intestine (612), Spleen (603), Testes (408), Thymus (1143), and Whole brain (1315) |

[2] https://github.com/pinellolab/scATAC-benchmarking/tree/master/Real_Data/Buenrostro_2018
[3] https://github.com/pinellolab/scATAC-benchmarking/tree/master/Real_Data/Cusanovich_2018_subset

**Table 2.** Pearson correlations between the simulated and real samples' bin means (excluding bins with a non-zero cell proportion $\geq 0.8$ in the real input data) and the non-zero cell proportion of bins, averaged across 20 simulation runs. Simulation is performed without including a Gaussian noise.

| | Correlation | |
|---|---|---|
| Cell type | Bin mean | Non-zero cell proportion |
| *Buenrostro2018* | | |
| GMP | 0.98 | 0.99 |
| CMP | 0.96 | 0.98 |
| pDC | 0.97 | 0.97 |
| HSC | 0.98 | 0.98 |
| LMPP | 0.96 | 0.97 |
| MPP | 0.93 | 0.95 |
| mono | 0.92 | 0.94 |
| MEP | 0.96 | 0.96 |
| CLP | 0.91 | 0.92 |
| *Cusanovich2018* | | |
| Lung | 0.97 | 0.98 |
| Thymus | 0.99 | 0.99 |
| Heart | 0.95 | 0.96 |
| Spleen | 0.98 | 0.98 |
| PreFrontalCortex | 0.96 | 0.97 |
| LargeIntestine | 0.93 | 0.93 |
| BoneMarrow | 0.94 | 0.95 |
| Liver | 0.98 | 0.98 |
| Cerebellum | 0.91 | 0.91 |
| SmallIntestine | 0.81 | 0.83 |
| Kidney | 0.97 | 0.97 |
| WholeBrain | 0.96 | 0.98 |
| Testes | 0.86 | 0.87 |
| *PBMCs* | | |
| Cell1 | 0.99 | 1 |
| Cell2 | 0.99 | 1 |
| Cell3 | 0.99 | 1 |
| Cell4 | 0.99 | 0.99 |
| Cell5 | 0.98 | 0.99 |
| Cell6 | 0.98 | 0.99 |
| Cell7 | 0.99 | 0.99 |
| Cell8 | 0.68 | 0.77 |

**Table 3.** Peak calling comparison of peak-by-cell matrices extracted by simATAC from the simulated bin-by-cell matrices (including 30000 peak bins), and peaks obtained from MACS2 and Genrich peak callers. For each pair of peak lists in the table, we report the average of regions having an overlap with the other list over the average of the peak lists' lengths. The number of called peaks are provided for each dataset and software in the format of tool(number of called peaks). The reported values for the simulated datasets are the average of 20 simulation runs.

| | Buenrostro2018 | | | Cusanovich2018 | | | PBMCs | | |
|---|---|---|---|---|---|---|---|---|---|
| | MACS2(91088) | Genrich(79444) | simulated(ours) | MACS2(85898) | Genrich(31154) | simulated(ours) | MACS2(86490) | Genrich(55386) | simulated(ours) |
| MACS2 | 1 | 0.81 | 0.53 | 1 | 0.50 | 0.52 | 1 | 0.71 | 0.54 |
| Genrich | 0.81 | 1 | 0.63 | 0.50 | 1 | 0.81 | 0.71 | 1 | 0.76 |
| simulated(ours) | 0.53 | 0.63 | 1 | 0.52 | 0.81 | 1 | 0.54 | 0.76 | 1 |

**Table 4.** Clustering evaluation results. The NMI, AMI, and ARI scores obtained from the Seurat's clustering algorithm on peak-by-cell matrices obtained from input bin-by-cell matrix for 30000 peak bins (SnapTools), simATAC bin-by-cell matrices for 30000 peak bins and Gaussian noise mean: -0.5 and std: 0.5 (average of 20 simulation runs), MACS2 and Genrich peak-by-cell matrices.

| | Buenrostro2018 | | | Cusanovich2018 | | | PBMCs | | |
|---|---|---|---|---|---|---|---|---|---|
| Tool | NMI | AMI | ARI | NMI | AMI | ARI | NMI | AMI | ARI |
| SnapTools | 0.54 | 0.57 | 0.37 | 0.54 | 0.58 | 0.34 | 0.52 | 0.56 | 0.39 |
| MACS2 | 0.57 | 0.63 | 0.43 | 0.55 | 0.60 | 0.40 | 0.51 | 0.58 | 0.36 |
| Genrich | 0.55 | 0.61 | 0.40 | 0.54 | 0.57 | 0.33 | 0.56 | 0.60 | 0.38 |
| simATAC | 0.63 | 0.63 | 0.36 | 0.65 | 0.68 | 0.31 | 0.61 | 0.68 | 0.45 |

**Table 5.** Pearson correlations between the simulated and real samples' bin means (excluding bins with a non-zero cell proportion $\geq$ 0.8 in the real input data) and the non-zero cell proportion of bins, averaged across 20 simulation runs. Simulation is performed with Gaussian noise parameters mean: -0.3, standard deviation: 0.3.

| | Correlation | |
|---|---|---|
| Cell type | Bin mean | Non-zero cell proportion |
| *Buenrostro2018* | | |
| CMP | 0.96 | 0.98 |
| GMP | 0.98 | 0.98 |
| pDC | 0.96 | 0.96 |
| mono | 0.92 | 0.92 |
| *Cusanovich2018* | | |
| Heart | 0.95 | 0.95 |
| Kidney | 0.97 | 0.97 |
| LargeIntestine | 0.92 | 0.92 |
| Liver | 0.97 | 0.97 |
| *PBMCs* | | |
| Cell1 | 0.99 | 0.99 |
| Cell3 | 0.99 | 0.99 |
| Cell5 | 0.98 | 0.99 |
| Cell7 | 0.98 | 0.99 |

**Table 6.** Pearson correlations between the simulated and real samples' bin means (excluding bins with a non-zero cell proportion ≥ 0.8 in the real input data) and the non-zero cell proportion of bins, averaged across 20 simulation runs. Simulation is performed with Gaussian noise parameters mean: -0.4, standard deviation: 0.4.

| | Correlation | |
|---|---|---|
| Cell type | Bin mean | Non-zero cell proportion |
| *Buenrostro2018* | | |
| CMP | 0.96 | 0.97 |
| GMP | 0.98 | 0.97 |
| pDC | 0.95 | 0.94 |
| mono | 0.9 | 0.88 |
| *Cusanovich2018* | | |
| Heart | 0.93 | 0.92 |
| Kidney | 0.96 | 0.95 |
| LargeIntestine | 0.9 | 0.89 |
| Liver | 0.96 | 0.96 |
| *PBMCs* | | |
| Cell1 | 0.99 | 0.99 |
| Cell3 | 0.99 | 0.99 |
| Cell5 | 0.98 | 0.98 |
| Cell7 | 0.98 | 0.98 |

**Table 7.** Clustering evaluation results. The NMI, AMI, and ARI scores obtained from the SnapATAC clustering algorithm. The metrics are the average of 20 simulation runs for each sparsity adjustment factor value.

| | Buenrostro2018 | | | Cusanovich2018 | | | PBMCs | | |
|---|---|---|---|---|---|---|---|---|---|
| Sparsity adjustment factor | NMI | AMI | ARI | NMI | AMI | ARI | NMI | AMI | ARI |
| 0.8 | 0.70 | 0.82 | 0.53 | 0.96 | 0.97 | 0.95 | 0.76 | 0.85 | 0.69 |
| 0.9 | 0.68 | 0.80 | 0.50 | 0.96 | 0.97 | 0.95 | 0.78 | 0.87 | 0.73 |
| 1 | 0.71 | 0.83 | 0.54 | 0.96 | 0.97 | 0.94 | 0.78 | 0.87 | 0.71 |
| 1.1 | 0.70 | 0.82 | 0.51 | 0.96 | 0.97 | 0.94 | 0.77 | 0.86 | 0.71 |
| 1.2 | 0.73 | 0.84 | 0.56 | 0.94 | 0.96 | 0.93 | 0.79 | 0.88 | 0.72 |

**Table 8.** simATAC estimation and simulation running time (in seconds) for three versions (V) of benchmark datasets' simulations, including Buenrostro2018, Cusanovich2018, and PBMCs. The running times are separately reported for estimation (Est) and simulation (Sim) steps for each cell group with specified simulated cell counts. Reported times are the average of three simulation runs with default input parameters.

| Cell type | Cell count | Est V1 | Est V2 | Est V3 | Sim V1 | Sim V2 | Sim V3 |
|---|---|---|---|---|---|---|---|
| Buenrostro2018(GMP) | 402 | 9.16 | 10.19 | 10.31 | 19.19 | 19.39 | 19.48 |
| Buenrostro2018(CMP) | 502 | 12.06 | 11.45 | 11.62 | 21.98 | 21.67 | 21.98 |
| Buenrostro2018(pDC) | 141 | 3.06 | 3.03 | 3.30 | 11.91 | 11.65 | 11.87 |
| Buenrostro2018(HSC) | 347 | 7.56 | 7.16 | 7.72 | 17.32 | 17.25 | 18.21 |
| Buenrostro2018(LMPP) | 160 | 3.29 | 3.38 | 3.34 | 12.33 | 12.16 | 12.28 |
| Buenrostro2018(MPP) | 142 | 3.10 | 3.08 | 3.09 | 11.98 | 11.53 | 11.68 |
| Buenrostro2018(mono) | 64 | 1.59 | 1.42 | 1.43 | 10.05 | 9.67 | 9.74 |
| Buenrostro2018(MEP) | 138 | 2.94 | 3.09 | 3.07 | 11.92 | 11.83 | 11.94 |
| Buenrostro2018(CLP) | 78 | 1.77 | 1.76 | 1.67 | 10.21 | 10.03 | 10.06 |
| Cusanovich2018(Lung) | 1499 | 35.76 | 34.79 | 35.04 | 40.91 | 41.42 | 41.14 |
| Cusanovich2018(Thymus) | 1143 | 26.54 | 26.56 | 26.53 | 33.26 | 33.27 | 33.24 |
| Cusanovich2018(Heart) | 1148 | 26.09 | 24.77 | 25.26 | 33.41 | 33.34 | 33.32 |
| Cusanovich2018(Spleen) | 603 | 13.17 | 12.13 | 12.43 | 20.71 | 20.52 | 20.57 |
| Cusanovich2018(PreFrontalCortex) | 894 | 20.44 | 19.59 | 19.72 | 27.39 | 27.55 | 27.67 |
| Cusanovich2018(LargeIntestine) | 1063 | 25.09 | 23.74 | 24.18 | 31.62 | 31.43 | 31.66 |
| Cusanovich2018(BoneMarrow) | 1261 | 31.70 | 29.64 | 29.88 | 35.92 | 35.57 | 35.75 |
| Cusanovich2018(Liver) | 925 | 20.67 | 19.97 | 20.37 | 29.38 | 28.31 | 28.46 |
| Cusanovich2018(Cerebellum) | 342 | 6.21 | 6.01 | 6.44 | 14.19 | 14.13 | 14.34 |
| Cusanovich2018(SmallIntestine) | 612 | 13.35 | 12.85 | 12.70 | 20.84 | 20.83 | 20.73 |
| Cusanovich2018(Kidney) | 965 | 21.79 | 21.07 | 21.25 | 29.34 | 28.91 | 28.86 |
| Cusanovich2018(WholeBrain) | 1315 | 30.69 | 29.69 | 30.00 | 37.10 | 37.37 | 36.95 |
| Cusanovich2018(Testes) | 408 | 7.91 | 7.46 | 8.20 | 16.00 | 15.99 | 16.09 |
| PBMCs(Cell1) | 963 | 24.59 | 23.42 | 23.79 | 34.13 | 34.29 | 33.81 |
| PBMCs(Cell2) | 939 | 23.94 | 22.83 | 23.15 | 33.17 | 33.30 | 33.02 |
| PBMCs(Cell3) | 928 | 24.07 | 22.66 | 23.14 | 33.23 | 33.41 | 33.01 |
| PBMCs(Cell4) | 789 | 20.48 | 18.97 | 19.37 | 29.85 | 29.53 | 29.26 |
| PBMCs(Cell5) | 634 | 15.40 | 14.32 | 14.77 | 25.20 | 25.40 | 25.17 |
| PBMCs(Cell6) | 619 | 15.14 | 14.30 | 14.58 | 25.11 | 24.71 | 24.49 |
| PBMCs(Cell7) | 362 | 7.67 | 7.73 | 7.91 | 17.58 | 17.45 | 17.39 |
| PBMCs(Cell8) | 101 | 2.60 | 2.49 | 2.55 | 11.21 | 10.84 | 10.79 |

**Table 9.** Table of abbreviations

| Abbreviation | Complete form |
|---|---|
| ATAC-seq | Assay for Transposase-Accessible Chromatin Sequencing |
| scATAC-seq | Single-Cell Assay for Transposase-Accessible Chromatin Sequencing |
| SCE | SingleCellExperiment |
| GMM | Gaussian Mixture Model |
| 10xG | 10x Genomics Chromium |
| PBMCs | Peripheral Blood Mononuclear Cells |
| MAD | Median Absolute Deviation |
| MAE | Mean Absolute Error |
| RMSE | Root Mean Square Error |
| NMI | Normalized Mutual Information |
| AMI | Adjusted Mutual Information |
| ARI | Adjusted Rand Index |
| sciATAC-seq | Single-Cell Combinatorial Indexing ATAC-seq |
| MI-ATAC | Multi-Index Single-Cell ATAC-seq |
| snATAC-seq | Single-Nucleus ATAC-seq |
| MI | Mutual Information |
| RI | Rand Index |

### 1 NOTE S1: PEAK-BY-CELL MATRIX GENERATION PIPELINE

Having raw BAM files of the benchmark datasets (one file per single-cell), we used the Picard tool (version 2.23.3) *MarkDuplicates* function to remove duplicate reads (https://broadinstitute.github.io/picard/). We used samtools (version 1.10) for filtering properly paired and uniquely mapped reads having a mapping quality $> 30$ (http://samtools.github.io).

We then run bedtools (version 2.27.1) *coverage* function to find the number of aligned reads in the obtained peaks (https://bedtools.readthedocs.io/en/latest/).

We run Genrich with -j, -r, -m 30, and -v input arguments (https://github.com/jsh58/Genrich), and MACS2 with -f BAMPE, –nomodel, and -g inputs (https://github.com/macs3-project/MACS), and generated peak-by-cell matrices.

#### 2 NOTE S2: CLUSTERING PERFORMANCE ASSESSMENT

Having the predicted and true labels obtained from Chen et al.[11], we assessed the clustering performance with three well-established metrics, including normalized mutual information (NMI), adjusted mutual information (AMI), and adjusted Rand index (ARI). Considering $gt$ and $pred$ as the ground truth and predicted labels, respectively, their entropy are defined by

$$H(gt) = -\sum_{i=1}^{|gt|} P(i)log(P(i)) \qquad \textbf{Equation } 1.$$

$$H(pred) = -\sum_{j=1}^{|pred|} P'(j)log(P'(j)) \qquad \textbf{Equation } 2.$$

where $P(i) = \frac{|gt_i|}{N}$ and $P'(j) = \frac{|pred_j|}{N}$, considering $N$ the number of cells, $gt_i$ the number of cells having label $i$ in $gt$, and $pred_j$ the number of cells having label $j$ in $pred$. The mutual information (MI) between $gt$ and $pred$ is calculated by

$$MI(gt, pred) = \sum_{i=1}^{|gt|} \sum_{j=1}^{|pred|} P(i,j)log(\frac{P(i,j)}{P(i)P'(j)}) \qquad \textbf{Equation } 3.$$

where $P(i,j) = \frac{|gt_i \cap pred_j|}{N}$. Having the MI, the NMI metric is defined as

$$NMI(gt, pred) = \frac{MI(gt, pred)}{max(H(gt), H(pred))} \qquad \textbf{Equation } 4.$$

Considering $E$ as the expected value, AMI is calculated by

$$AMI(gt, pred) = \frac{MI - E[MI]}{mean(H(gt), H(pred)) - E[MI]} \qquad \textbf{Equation } 5.$$

We also obtained ARI metric from

$$ARI(gt, pred) = \frac{RI - E[RI]}{max(RI) - E[RI]} \qquad \textbf{Equation } 6.$$

where the Rand Index (RI) computes a similarity measure between two clusterings by considering all pairs of samples and counting pairs that are assigned in the same or different clusters in the predicted and true clusterings [4].

[4] https://scikit-learn.org/stable/modules/clustering.html

#### REFERENCES


1. Alicia N. Schep, Beijing Wu, Jason D. Buenrostro, and William J. Greenleaf. ChromVAR: Inferring transcription-factor-associated accessibility from single-cell epigenomic data. *Nature Methods*, 2017.
2. M. Ryan Corces, Jason D. Buenrostro, Beijing Wu, Peyton G. Greenside, Steven M. Chan, Julie L. Koenig, Michael P. Snyder, Jonathan K. Pritchard, Anshul Kundaje, William J. Greenleaf, Ravindra Majeti, and Howard Y. Chang. Lineage-specific and single-cell chromatin accessibility charts human hematopoiesis and leukemia evolution. *Nature Genetics*, 2016.
3. Jason D. Buenrostro, Beijing Wu, Ulrike M. Litzenburger, Dave Ruff, Michael L. Gonzales, Michael P. Snyder, Howard Y. Chang, and William J. Greenleaf. Single-cell chromatin accessibility reveals principles of regulatory variation. *Nature*, 2015.
4. Darren A. Cusanovich, Riza Daza, Andrew Adey, Hannah A. Pliner, Lena Christiansen, Kevin L. Gunderson, Frank J. Steemers, Cole Trapnell, and Jay Shendure. Multiplex single-cell profiling of chromatin accessibility by combinatorial cellular indexing. *Science*, 2015.
5. Xingqi Chen, Ulrike M. Litzenburger, Yuning Wei, Alicia N. Schep, Edward L. LaGory, Hani Choudhry, Amato J. Giaccia, William J. Greenleaf, and Howard Y. Chang. Joint single-cell DNA accessibility and protein epitope profiling reveals environmental regulation of epigenomic heterogeneity. *Nature Communications*, 2018.
6. Sebastian Preissl, Rongxin Fang, Hui Huang, Yuan Zhao, Ramya Raviram, David U. Gorkin, Yanxiao Zhang, Brandon C. Sos, Veena Afzal, Diane E. Dickel, Samantha Kuan, Axel Visel, Len A. Pennacchio, Kun Zhang, and Bing Ren. Single-nucleus analysis of accessible chromatin in developing mouse forebrain reveals cell-type-specific transcriptional regulation. *Nature Neuroscience*, 2018.
7. Ansuman T Satpathy, Jeffrey M Granja, Kathryn E Yost, Yanyan Qi, Francesca Meschi, Geoffrey P. McDermott, Brett N. Olsen, Maxwell R. Mumbach, Sarah E Pierce, M Ryan Corces, Preyas Shah, Jason C Bell, Darisha Jhutty, Corey M. Nemec, Jean Wang, Li Wang, Yifeng Yin, Paul G. Giresi, Anne Lynn S. Chang, Grace X Y Zheng, William J Greenleaf, and Howard Y Chang. Massively parallel single-cell chromatin landscapes of human immune cell development and intratumoral T cell exhaustion. *Nature Biotechnology*, 37(8):925–936, 2019.
8. Jason D. Buenrostro, M. Ryan Corces, Caleb A. Lareau, Beijing Wu, Alicia N. Schep, Martin J. Aryee, Ravindra Majeti, Howard Y. Chang, and William J. Greenleaf. Integrated Single-Cell Analysis Maps the Continuous Regulatory Landscape of Human Hematopoietic Differentiation. *Cell*, 2018.
9. Lei Xiong, Kui Xu, Kang Tian, Yanqiu Shao, Lei Tang, Ge Gao, Michael Zhang, Tao Jiang, and Qiangfeng Cliff Zhang. SCALE method for single-cell ATAC-seq analysis via latent feature extraction. *Nature Communications*, 2019.
10. Darren A. Cusanovich, Andrew J. Hill, Delasa Aghamirzaie, Riza M. Daza, Hannah A. Pliner, Joel B. Berletch, Galina N. Filippova, Xingfan Huang, Lena Christiansen, William S. DeWitt, Choli Lee, Samuel G. Regalado, David F. Read, Frank J. Steemers, Christine M. Disteche, Cole Trapnell, and Jay Shendure. A Single-Cell Atlas of In Vivo Mammalian Chromatin Accessibility. *Cell*, 2018.
11. Huidong Chen, Caleb Lareau, Tommaso Andreani, Michael E Vinyard, Sara P Garcia, Kendell Clement, Miguel A Andrade-Navarro, Jason D Buenrostro, and Luca Pinello. Assessment of computational methods for the analysis of single-cell ATAC-seq data, 2019.
